# Dynamic and reversible remapping of network representations in an unchanging environment

**DOI:** 10.1101/2020.10.05.326942

**Authors:** Isabel I. C. Low, Alex H. Williams, Malcolm G. Campbell, Scott W. Linderman, Lisa M. Giocomo

## Abstract

In response to environmental changes, the medial entorhinal cortex alters its single-cell firing properties. This flexibility in neural coding is hypothesized to support navigation and memory by dividing sensory experience into unique contextual episodes. However, it is unknown how the entorhinal circuit transitions between different representations, particularly when sensory information is not delineated into discrete contexts. Here, we describe spontaneous and abrupt transitions between multiple spatial maps of an unchanging task and environment. These remapping events were synchronized across hundreds of medial entorhinal neurons and correlated with changes in running speed. While remapping altered spatial coding in individual neurons, we show that features of the environment were statistically preserved at the population-level, enabling simple decoding strategies. These findings provoke a reconsideration of how medial entorhinal cortex dynamically represents space and broadly suggest a remarkable capacity for higher-order cortical circuits to rapidly and substantially reorganize their neural representations.

## Introduction

As animals engage in complex behaviors, they must dynamically integrate the sensory features of their environment with internal factors such as behavioral state. For example, arousal (Hulse et al., 2017; Salay et al., 2018; Vinck et al., 2015), satiety (Jennings et al., 2019), attention (Kentros et al., 2004), and running speed (Hardcastle et al., 2017; Hulse et al., 2017; Niell and Stryker, 2010; Vinck et al., 2015) have widespread impacts on cortical processing (Stringer et al., 2019) and influence how an animal interacts with its environment (Calhoun et al., 2019; Salay et al., 2018). In this way, the same set of sensory features associated with a given environment can drive unique neural representations when combined with different internal factors. However, this dynamic interaction between internal and external factors also presents a challenge to integration centers of the brain, which must balance reliable representations of sensory features with flexible responses to changing internal state. For example, during navigation animals encounter a continuous stream of sensory features while simultaneously experiencing bevarioral state changes. To encode unique episodes or contexts, the brain must integrate these internal and external factors in order to generate distinct neural representations that are consistent with the animal’s experience of the world.

A key neural substrate hypothesized to support this process is the medial entorhinal cortex (MEC), which contains neurons encoding the animal’s spatial position and orientation relative to features in the external world, such as environmental boundaries and objects (Diehl et al., 2017; Gil et al., 2018; Hafting et al., 2005; Høydal et al., 2019; Sargolini et al., 2006; Solstad et al., 2008). MEC position and orientation cells can alter their firing rates and their firing fields can rotate, shift, or switch locations in response to changes in the geometry, sensory features, or task demands associated with the environment—a phenomenon known as remapping (Barry et al., 2007; Boccara et al., 2019; Butler et al., 2019; Diehl et al., 2017; Fyhn et al., 2007; Keene et al., 2016; Krupic et al., 2015; Marozzi et al., 2015; Munn et al., 2020; Solstad et al., 2008). Previous studies of MEC remapping have primarily focused on how the firing fields of individual functionally-defined classes of cells (e.g. grid, border, or head direction cells) respond to distinct sets of environmental features. However, many MEC neurons do not fall into discrete functionally-defined classes (Hardcastle et al., 2017; Hinman et al., 2016), and theoretical models of MEC propose that neural dynamics emerge from interconnected networks of neurons (Burak and Fiete, 2009; Couey et al., 2013; Fuhs and Touretzky, 2006; McNaughton et al., 2006; Ocko et al., 2018; Pastoll et al., 2013). How MEC as a whole transitions between contextual representations, particularly when presented with environmental features that are not explicitly divided into distinct sets, is incompletely understood.

Here, we investigate how large populations of MEC neurons transition between spatial representations in an invariant external context. To do so, we use silicon probes to simultaneously record from hundreds of MEC neurons while mice navigate a tightly controlled virtual reality environment. We find that remapping events occur synchronously across the MEC population and can occur without any change in environmental features or task demands. Further, we demonstrate that each map corresponds to a single attractor manifold and that running speed correlates with neural variability, driving transitions between manifolds. Together, our findings bridge the gap between previous studies of flexibility in MEC coding (Barry et al., 2007; Boccara et al., 2019; Butler et al., 2019; Diehl et al., 2017; Fyhn et al., 2007; Keene et al., 2016; Krupic et al., 2015; Marozzi et al., 2015; Munn et al., 2020; Solstad et al., 2008) and existing theoretical models of MEC population dynamics (Burak and Fiete, 2009; Couey et al., 2013; Fuhs and Touretzky, 2006; McNaughton et al., 2006; Ocko et al., 2018; Pastoll et al., 2013).

## Results

### Spatial representations remap in an invariant virtual environment

We implemented a virtual reality (VR) navigation task in which head-fixed mice foraged for water rewards along an infinite track with landmark cues that repeated every 400 cm (Campbell et al., 2020)(fig. 1a, c). 6 mice experienced only a cue rich track (5 landmarks, fig. 1c, top; cool colors indicate mouse ID in all figures, fig. 1b, top)(n = 21 sessions, i.e. “cue rich, single-track” sessions), 6 mice experienced only a cue poor track (2 landmarks, fig. 1c, bottom; warm colors in all figures, fig. 1b, middle)(n = 11 sessions, i.e. “cue poor, single-track” sessions), and 5 mice experienced alternating blocks of cue rich and poor trials within each each session (n = 13 sessions, i.e. “double-track” sessions; purples in all figures, fig. 1b, bottom)(see also fig. S1). Mice could lick to request water in a visually marked reward zone, which appeared at random locations along the track. Mice consistently slowed (mean difference in running speed within vs. outside of reward zone ± standard error of the mean (SEM): 9.1 ± 1.5 cm/s; Wilcoxon two-sided signed-rank test, p = 3.9×10^−7^; n = 45 sessions in 17 mice) and licked for water (mean difference in lick number within vs. outside of zone ± SEM: 7.4 ± 0.3; Wilcoxon two-sided signed-rank test, p < 0.0001; n = 10,953 reward trials across 45 sessions in 17 mice) in reward zones, demonstrating familiarity with the task (fig. 1c-f). As reward locations were random, the task produced a spatially uniform distribution of running speeds (fig. 1g).

**Figure 1:**
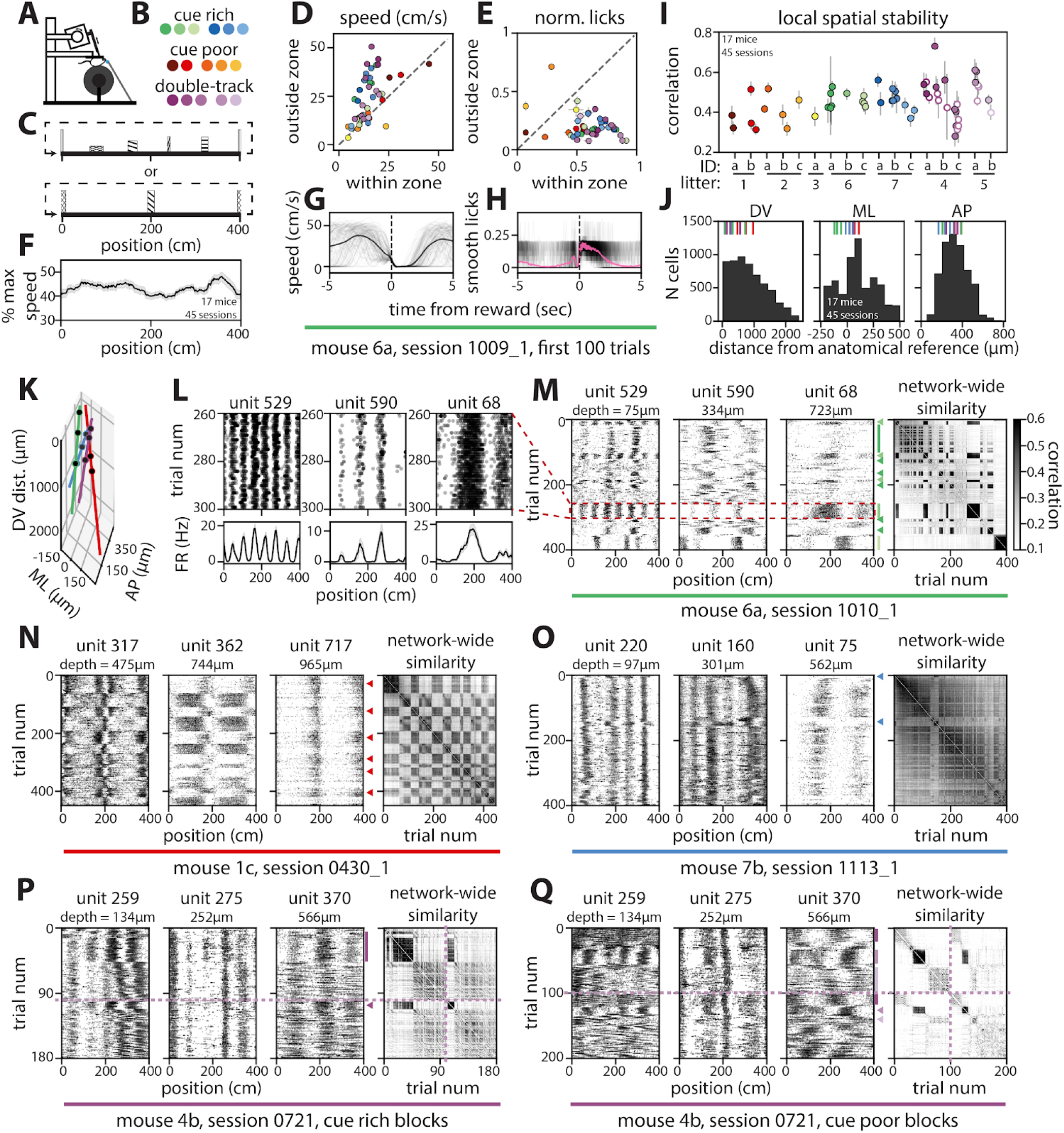
Spatial representations remap in an invariant virtual environment. (A) Schematic of recording set-up. (B) Color scheme for each track condition (colors represent individual mice and are maintained throughout this figure). (C) Side-view schematic of cue rich (top) and cue poor (bottom) environments. (D) Average running speed within versus outside of reward zones (points represent individual sessions). (E) Average fraction of licks that occurred within versus outside of the reward zones on each trial for each session. (F) Average running speed for each track position across all sessions (gray shading, SEM). (G) Running speed near rewards for the first 100 trials of an example session (gray traces, each trial; black line, average). (H) As in (G), but for smoothed lick count (pink line, average). (I) Average spatial correlation of nearby trials for each session. Double-track sessions (purple) show cue rich (filled) and cue poor (open) blocks separately. (J) Locations for all recorded MEC units relative to the dorsal border of MEC (dorsal-ventral (DV) = 0), middle of MEC (medial-lateral (ML) = 0), and back of brain (anterior-posterior (AP) = 0)(hash-marks correspond to example units in K-Q). (K) Probe locations for examples in (L-Q)(lines, probe; points, example units) relative to anatomical boundaries as in (J). (L) Raster plots (top) and tuning curves (bottom) for three example cells from one session (black lines, average firing rate; shading, SEM). (M-O) Rasters of example units (left; arrowheads and lines indicate distinct maps) and network-wide trial-by-trial similarity matrices (right) from 3 example single-track sessions (M, n = 142 cells; N, n = 227 cells; O, n = 139 cells). (P, Q) As in (M-O), but for one double-track session split into cue rich (P) and cue poor (Q) trial blocks (dashed lines indicate breaks between blocks; n = 55 cells). (C, D, H, I) N = 5,963 cells in 45 sessions across 17 mice. (See also Fig. S1, S2, S3, S4.)(Methods). This procedure resulted in a trial-by-trial similarity matrix of network-wide spatial representations for each session (fig. 1m-q right; fig. S3). Sessions were on average locally spatially stable (mean moving average correlation of 5 nearest trials ± SEM: 0.455 ± 0.011; n = 32 single-track sessions and 26 cue rich or poor blocks from 13 double-track sessions in 17 mice)

To record neural activity as mice traversed the VR tracks, we acutely inserted Neuropixels silicon probes (Jun et al., 2017) into MEC in up to six recording sessions per mouse (up to three sessions per hemisphere), each associated with a unique probe insertion (fig. 1a, k; fig. S1). We obtained simultaneous recordings from hundreds of MEC neurons in individual mice across a large portion of the dorsal to ventral axis of MEC (n = 5,963 cells across 45 sessions in 17 mice)(fig. 1j, k; fig. S1). Many individual neurons exhibited spatially periodic firing patterns, with decreasing spatial frequencies along the dorsal to ventral axis of MEC (fig. 1l; fig. S2), consistent with known grid cell properties (Brun et al., 2008; Fyhn et al., 2008; Hafting et al., 2005). To characterize spatial coding across all co-recorded neurons, we estimated each neuron’s position-aligned firing rate on each trial and computed the correlation between all spatial representations for each pair of trials (Methods), though in some cases neural representations appeared untethered from landmarks for part or all of the session (mean spatial correlation range: 0.28 to 0.73; within session/block interquartile range min to max: 0.039 to 0.394)(fig. 1i; fig. S2). Nonetheless, many neurons still exhibited spatially periodic firing in these unstable trial blocks (fig. S2).

In many recording sessions, we observed clear changes in the spatial firing patterns of single neurons distributed across MEC (i.e., remapping events), as well as in network-wide trial-by-trial similarity matrices (fig. 1m-q, arrowheads and lines; fig. S3). Unlike previous reports of remapping in MEC (Barry et al., 2007; Boccara et al., 2019; Butler et al., 2019; Diehl et al., 2017; Fyhn et al., 2007; Keene et al., 2016; Krupic et al., 2015; Marozzi et al., 2015; Munn et al., 2020; Solstad et al., 2008), these remapping events occurred without any change to environmental sensory cues or task demands. Importantly, remapping events did not reflect shifts in the location of the recording probe, as spike waveforms remained stable across remapping events (fig. S4), we observed similar remapping events in recordings using tetrodes (fig. S4), and adjacent brain regions did not show comparable remapping in this task (fig. S1). In many recordings, MEC cells switched abruptly between stable spatial representations (i.e. maps), with cells returning repeatedly to one of several distinct maps within a single session, resulting in a checkerboard pattern in the trial-by-trial similarity matrices (fig. 1m-q; fig. S1, S3). In other sessions, spatial representations underwent a single transition between stable maps (fig. S3) or transitioned abruptly between spatially stable and unstable coding regimes (fig. 1m, q; fig. S2). While the frequency and stability of remapping was thus heterogeneous across mice and sessions, in all cases remap events appeared to recruit co-recorded neurons all along the dorsal to ventral MEC axis (fig. 1m-q, right; fig. S1-3, S5).

### Entorhinal neurons reversibly remap between different spatial representations

To group trials with similar network-wide spatial activity, we applied k-means clustering to each session (fig. 2a). The k-means model assigns a single cluster label to each trial and these cluster labels often visibly matched the checkerboard pattern in trial-by-trial similarity matrices (fig. 2b; fig. S3, S5). Despite making the strict assumption that each trial belongs to a single spatial map, a 2-cluster k-means model consistently approached the performance of a less constrained uncentered PCA model, which allows each trial to contain a blend of multiple spatial maps (fig. 2c, d; performance is measured by uncentered *R*^2^). These results suggest that remapping events were well-approximated by discrete transitions between spatial maps. Using the relative performance of k-means to PCA, we identified 18/32 single-track sessions across 8 mice that were adequately fit by a 2-cluster k-means model (fig. 2e, green points; performance gap with PCA < 70% relative to shuffle, 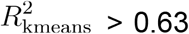). Of these 18 sessions, 10 exhibited three or more remap events (n = 4 mice), thus visiting each of the 2 maps at least twice (mean remap events ± SEM: 6.2 ± 1.8; range: 1 to 27; n = 18 sessions across 8 mice)(fig. S5). We focused subsequent analysis on these 18 “2-map sessions,” as they were the simplest and most common case (example sessions with periods of unstable spatial coding are shown in fig. S2; example sessions with > 2 spatial maps, fig. S5).

**Figure 2:**
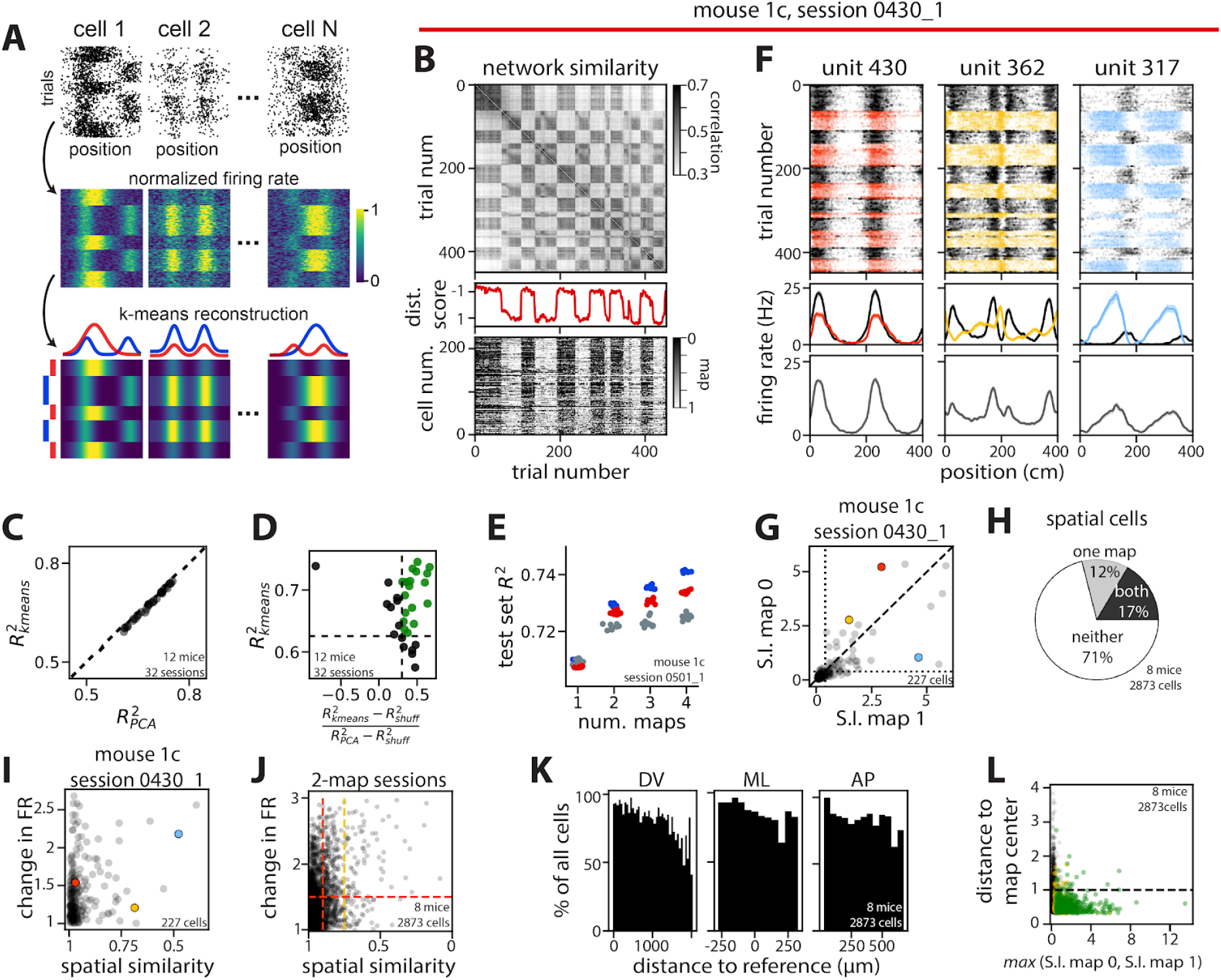
Entorhinal neurons reversibly remap between different spatial representations. (A) Schematic of k-means clustering strategy showing conversion of raw spikes (top) to normalized firing rate (center) and resulting low-dimensional k-means clustering estimate of the neural activity (bottom). (B) Trial-by-trial similarity (top, as in fig. 1M-O), distance score for population-wide neural activity by trial (middle; score = 1, in map 1 centroid; score = −1, map 0), and single neuron distance to k-means cluster centroid across trials, sorted by depth (bottom; gray, midway between maps; black, at or beyond map 0 centroid; white, map 1) for an example session. (C) 2-factor k-means versus 2-factor PCA performance for all single-track sessions (n = 32 sessions, 12 mice). (D) Selection criteria for 2-map sessions (green points)(n = 32 sessions, 12 mice). (E) Model performance for uncentered PCA (blue), k-means (red), and k-means on shuffled data (gray) for an example session. (F) Single-neuron spiking (top) and tuning curves (middle) for example cells from an example session, colored/divided by k-means cluster labels (black, map 0; color, map 1), versus averaged over the full session (bottom)(solid line, trial-averaged firing rate; shading, SEM; color scheme denotes cell identity and is preserved in G, I, L). (G) Spatial information (S.I.) for single neurons in map 0 versus map 1 for an example session. (H) Percent of MEC neurons from 2-map sessions that were spatial in just one map (gray), both maps (black), or neither map (white). (I) Absolute fold change in firing rate versus cosine similarity of single-map tuning curves for neurons from an example session. (J) As in (I), but for all neurons from 2-map sessions (red vertical dashes, 75% spatial similarity; yellow dashes, 90% similarity; 50% change in firing rate). (K) Percent of all cells that were consistent remappers by location in MEC. (L) Distance to k-means cluster centroid by highest average S.I. across maps (black dashes, consistent remapper threshold; green, spatial in both maps; gold spatial in one map). N = 2873 cells, 18 sessions, 8 mice, unless noted. (See also Fig. S3, S5.)

In 2-map sessions, many single neurons exhibited distinct position tuning curves within each k-means identified spatial map (fig. 2f, middle row; see also fig. S2, S5). Averaging neural activity over the entire session—as is typically done in single-cell analyses of MEC coding—obscured these differences in tuning (fig. 2f, bottom row). Further, we observed changes in spatial information across the two maps (mean absolute change in spatial information ± SEM: 55.5 ± 1.4%; n = 2,873 cells in 8 mice)(fig. 2g), such that only a small percentage of all cells were significantly spatial in both maps (mean ± SEM: spatial in both maps = 17 ± 2%; one map = 13 ± 1%; neither map = 71 ± 3%; p < 0.05; n = 18 sessions in 8 mice)(fig. 2h). We further quantified these differences by calculating the change in peak firing rate (i.e. rate remapping) and in spatial similarity (i.e. global remapping) across the two maps for each cell (Knierim et al., 1998; Muller and Kubie, 1987; O’Keefe and Conway, 1978) (Methods). Across all cells (spatial and non-spatial), we observed rate remapping (fig. 2f, left), global remapping (fig. 2f, middle), and combinations of both (n = 2,873 cells in 8 mice)(fig. 2f right, fig. 2i, j). Rate remapping was common (818 out of 2,873 cells > 1.5-fold change in firing rate)(fig. 2j red dashes, horizontal). Large differences in spatial representations across maps were rare (118 cells < 75% tuning curve similarity)(fig. 2j yellow dashes), but many cells exhibited some change in spatial preference (568 cells < 90% similarity)(fig. 2j, red dashes, vertical). Altogether, 300 cells showed both rate and global remapping (> 1.5-fold change in firing rate, < 90% spatial similarity)(fig. 2i, j, red dashes). Further, the majority of single cells throughout MEC changed their firing properties in precise agreement with the population-derived remapping events (i.e. “consistent remappers,” 2,405 out of 2,873 cells distance to cluster centroid < 1)(fig. 2b bottom, k, l; fig. S4)(Methods). There was some regional variability in the proportion of these cells (fig. 2k), but nearly all cells that were significantly spatial in at least one map were consistent remappers (789 out of 830 spatial cells)(fig. 2l, green and gold points). Together, these results indicate that remapping events recruit a large portion of the MEC neural population and reflect changes in the spatial coding properties of this circuit.

### Positional information is conserved at a population level across distinct entorhinal spatial maps

MEC neurons project to multiple brain areas involved in spatial information processing, including the hippocampus, retrosplenial cortex, and the subiculum (Kerr et al., 2007). How might these areas make use of positional information encoded in MEC, in spite of network-level remapping? To investigate this question, we fit circular-linear regression models to predict position from neural activity (fig. 3a-c) and found that these simple decoding models were robust to remapping. Performance was comparable between models trained and tested on trials from a single map, and models trained and tested on trials from both maps (fig. 3d, e; mean score ± SEM: train/test map 0 = 0.60 ± 0.06; train/test map 1 = 0.65 ± 0.05; train/test both maps = 0.65 ± 0.06; Kruskal-Wallis H-test, p = 0.87; n = 18 sessions in 8 mice). Performance significantly worsened, however, when training data and testing data were taken from distinct maps (fig. 3d, f, g; Wilcoxon two-sided signed-rank test, p = 1.68−10^−7^; n = 36 model pairs). These findings indicate that the two maps are distinct from one another, but given training data from both maps it is possible to find a decoder that is robust to remapping. Intuitively, this can be accomplished when a decoder selectively makes use of neurons with only minor remapping, or, more generally, when remapping occurs in the null space of the decoder weights (Kaufman et al., 2014; Rule et al., 2020). Altogether, these results indicate that, in principle, downstream brain areas can still exploit positional information encoded in MEC in the presence of remapping.

**Figure 3:**
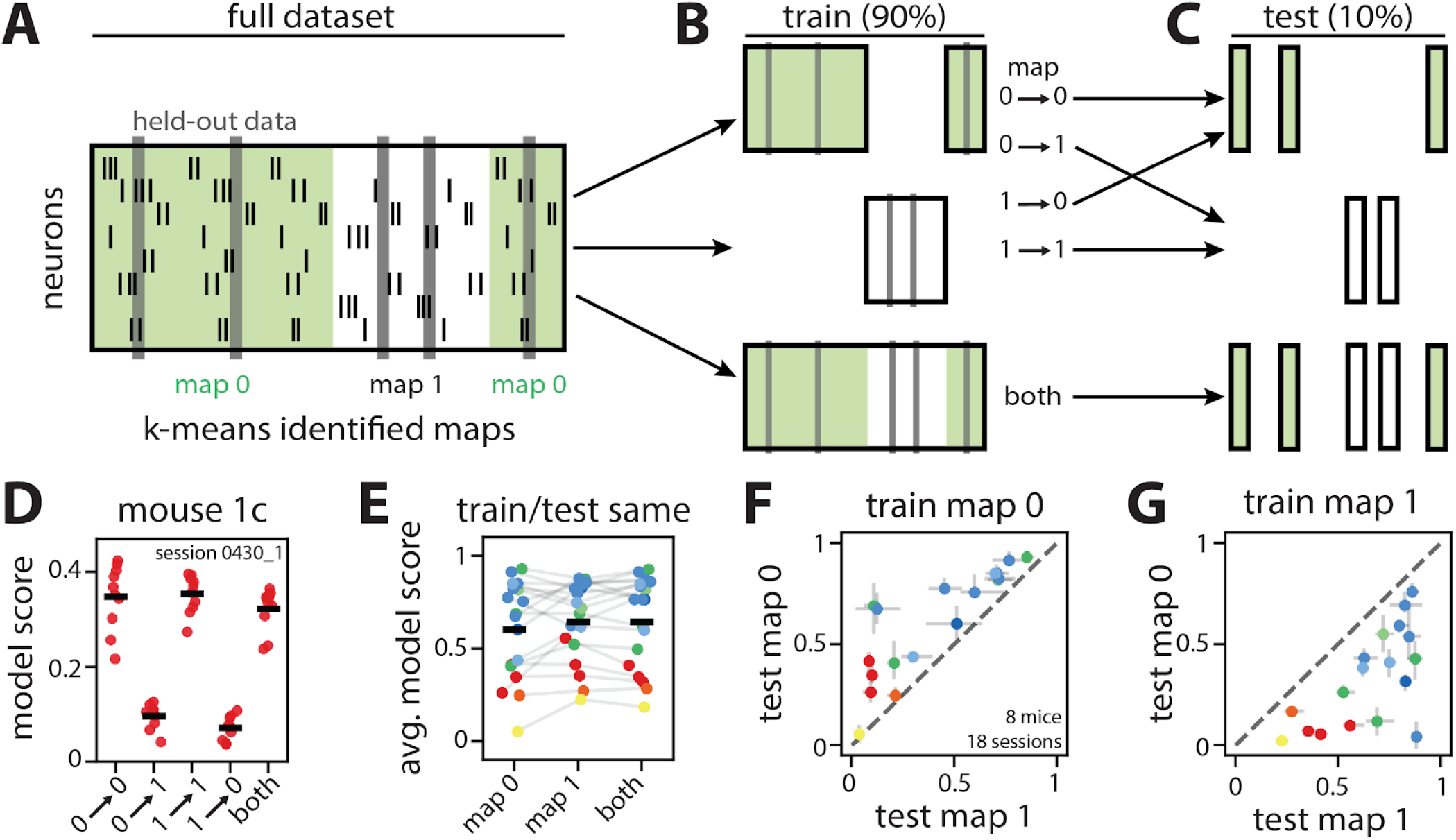
Positional information is conserved at a population level across distinct entorhinal spatial maps. (A-C) Schematic of decoder training and testing strategy (green shading, map 0; white, map 1). (A) Population-wide spiking activity (hash marks) was divided into k-means clusters and 10% of data was held out for testing (gray shading). (B) Decoders were trained on spiking data from either map 0 (top), map 1 (middle), or both maps (bottom). (C) Each model was then used to predict position using held-out spiking data from either map 0 (top), map 1 (middle), or both maps (bottom). Note that models trained only on data from map 0 or map 1 were also used to predict position using only data from the other map (diagonal arrows from B to C; labels indicate test map→train map). (D) Decoder performance using training data from each map alone (test→train) or from both maps (“both”) for an example session (score = 0, chance; score = 1, perfect prediction; N = 227 cells). (E) Decoder performance for models trained and tested on data from the same map (map 0, map 1) or from both maps across all 2-map sessions. (F, G) Decoder performance within versus across maps for all sessions (Grey bars, interquartile range). (E-G) Points indicate individual sessions; colors, mouse identity. N = 2873 cells, 18 sessions, 8 mice, unless noted.

### Neural activity transitions between geometrically aligned ring attractor manifolds

MEC representations of space thus display a capacity for flexibility (e.g. spontaneous remapping), as well as reliability (e.g. consistent decoding performance). To reconcile these two aspects of the circuit, we characterized the geometry of position coding in N-dimensional firing rate space (where N denotes the number of simultaneously recorded neurons). For the continuous 1D virtual environments used in this study, attractor network models (Burak and Fiete, 2009; Fuhs and Touretzky, 2006; Guanella et al., 2007) predict that the trajectory of network activity should trace out a 1D ring manifold as the animal moves through space (fig. 4a). Each k-means cluster centroid provides an empirical estimate of this attractor manifold and, indeed, the low-dimensional linear embedding of each centroid had a qualitative ring structure (fig. 4b-c). Manifolds derived from cue poor environments were tightly wound around themselves, as quantified by an entanglement metric (fig. 4d), indicating that there may be limited discriminability between the first and second halves of the track in cue poor environments. These results suggest that the number of landmark cues alters the geometry of the spatial map without altering its topology as a 1D ring.

**Figure 4:**
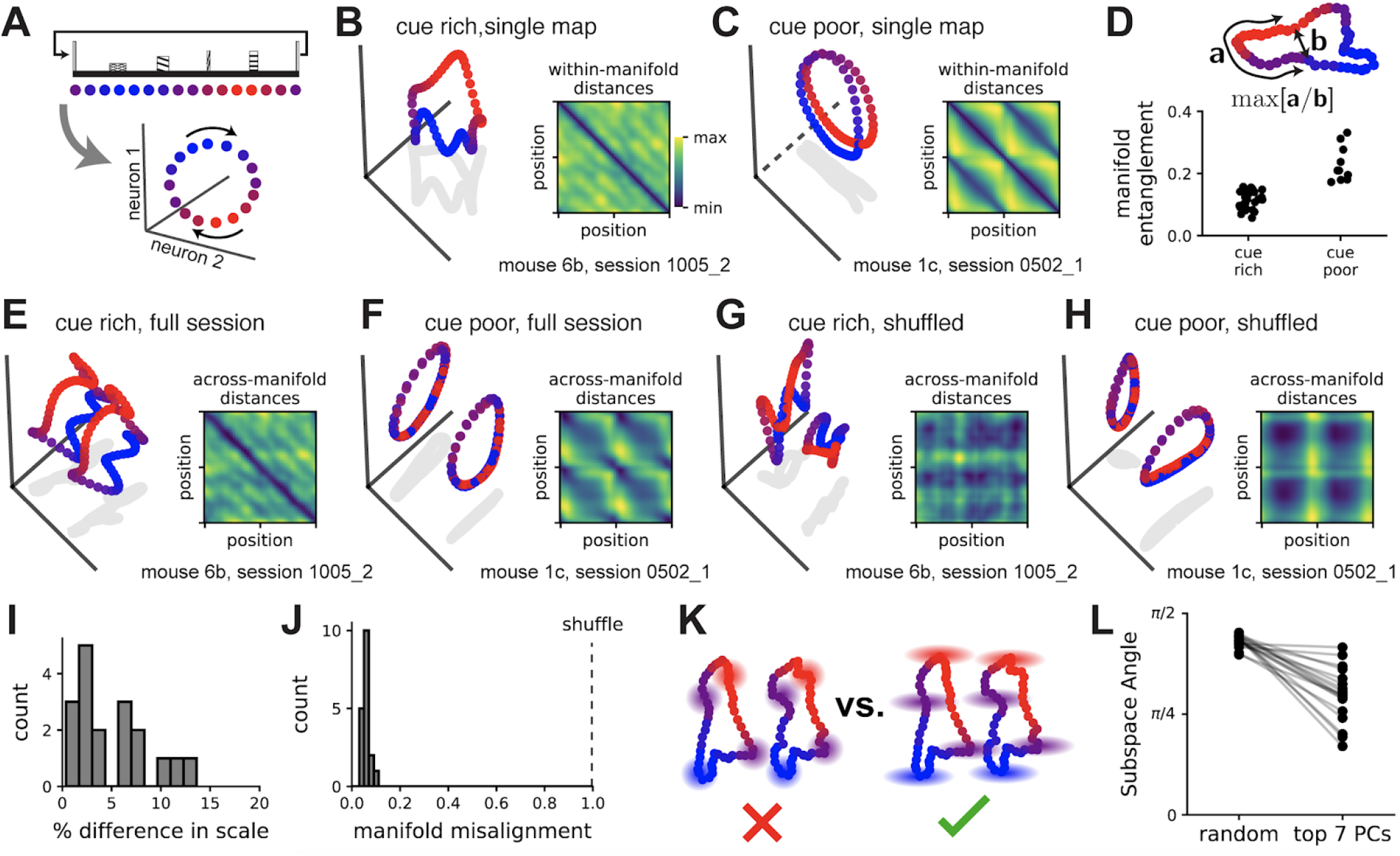
Neural activity transitions between geometrically aligned ring attractor manifolds. (A) Schematic: we expect the 1D environment (top, side-view shown) to produce a circular 1D trajectory (i.e., ring manifold) in neural firing rates (bottom; color scheme indicates position and is preserved throughout this figure). (B) PCA projection of a single map (k-means centroid) from a cue rich recording session (n = 149 cells). (Inset) Pairwise distances in neural firing rates across all points (i.e. spatial position bins) in the manifold (color code blue, minimum value; yellow, maximum). (C) As in (B), but for a cue poor recording session (n = 227 cells). The linear projection uses two principal components and a third orthogonal axis (dashed line) that maximally distinguishes between the first and second half of the track. (D) Manifold entanglement in cue rich and cue poor environments (top: schematic of entanglement measure; n = 10 cue poor manifolds, n = 26 cue rich manifolds). (E) PCA projection of two manifolds extracted from a 2-map cue rich session. (Inset) Across-manifold distances in neural firing rates for every pair of points (color code as in B, C insets). (F) As in (E), but for a cue poor session. (G, H) As in (E, F), respectively, but after applying a random rotation to each manifold. (I) Percent difference in manifold sizes for spatial maps recorded in the same behavioral session (n = 18 sessions). (J) Manifold misalignment scores, normalized to range between zero and one (n = 18 sessions). (K) Schematics showing that variability around manifolds could be isotropically distributed (left) or magnified in the direction of the other manifold (right). (L) Angles between the remapping dimension and the subspace spanned by the top 7 principal components, compared to angles formed with random 7D subspaces (averaged across 1,000 samples). All sessions (n = 18) displayed smaller angles than their shuffle control (Wilcoxon rank-sum test, p < 1e-6). (See also Fig. S6.)

We next used PCA to simultaneously visualize the two manifolds of each 2-map session in the same low-dimensional space. In both cue rich and cue poor environments, the manifolds appeared geometrically aligned such that position coding was translated along a single dimension in neural activity space (fig. 4e-f); applying a random rotation to each manifold qualitatively disrupted this alignment (fig. 4g-h). Using standard tools from statistical shape analysis (Methods), we found the optimal rescaling, rotation, and reflection to align the manifolds in each session. The two manifolds required only modest rescaling (fig. 4i, 11/17 sessions ≤ 10% difference in scale), suggesting minor changes in overall population firing rate. Further, in all cases, the observed manifolds were within 7% of the optimal rotational alignment, measured relative to the root-mean-squared-error under random rotations and reflections (fig. 4j), suggesting that spatial tuning was largely preserved at the population level (fig. 2i, j). Thus, remapping largely corresponded to a translation in neural representations. This observation is not a necessary consequence of existing theoretical frameworks—when we developed a simple extension of the classic ring attractor circuit to support bistable ring manifolds, this model could not account for the geometrical alignment observed experimentally (Supplementary Text; fig. S6a-g). This suggests that the alignment of the spatial manifolds may be a functionally important feature of the circuit. Indeed, if remapping is well-described by a translation along a single dimension, any decoder that is insensitive to this dimension will be robust to remapping, providing a simple strategy by which a downstream region could decode position. In the absence of manifold alignment, more complex decoders would be required (Supplementary Text; fig. S6h-n).

As the k-means-identified ring manifolds show only the average neural activity within each map, we next asked how single-trial variability around these manifolds was structured. Applying PCA to the residuals of the k-means model showed that the remaining variance was preferentially oriented in the dimension separating the two manifolds (fig. 4k-l). Importantly, the scale of this variability was limited so that the two manifolds often remained well-separated (i.e. it is generally appropriate to conceptualize the maps as two separate ring manifolds, rather than a single hollow cylinder). This result suggests that variability in network activity could predispose the network to remap (i.e. to jump from one ring manifold to the other), with attractor dynamics locally stabilizing the network activity within each map. The geometrical alignment of the two manifolds supports this hypothesis, as it implies that simple perturbations could induce remapping without introducing errors into the MEC network positional estimate (Supplementary Text; fig. S6h-n).

### Remapping events and neural variability correlate with slower running speeds

As the task and environment in our experiments did not change within a given session, we next examined whether the single-trial variability correlated with aspects of the animal’s behavior. For example, the k-means clustering model does not account for running speed, but speed is known to modulate spatial representations in MEC (Bant et al., 2020; Hardcastle et al., 2017). First, we asked whether the animal’s running speed was different on “remap trials” (defined as the two trials book-ending each transition from one map to the other), compared to the intervening “stable blocks” (defined as the block of trials at least two trials away from a remap event, which all reside in the same map)(fig. 5a, c). We restricted our analysis to 2-map sessions that contained at least three remap events and to stable blocks of at least five trials (n = 10 sessions in 4 mice; see also fig. S1, S6 for analysis of double-track sessions and >2-map sessions, respectively)(Methods). Across most of these sessions, the animal’s average running speed on remap trials was lower compared to its average running speed in the preceding stable block (7/10 sessions; speeds were equal in 3 sessions; mean percent difference in running speed ± SEM: 12.7 ± 2.4%; Wilcoxon two-sided signed-rank test, p = 1.06×10^−5^; n = 103 remap trial/stable block pairs)(fig. 5a-e).

**Figure 5:**
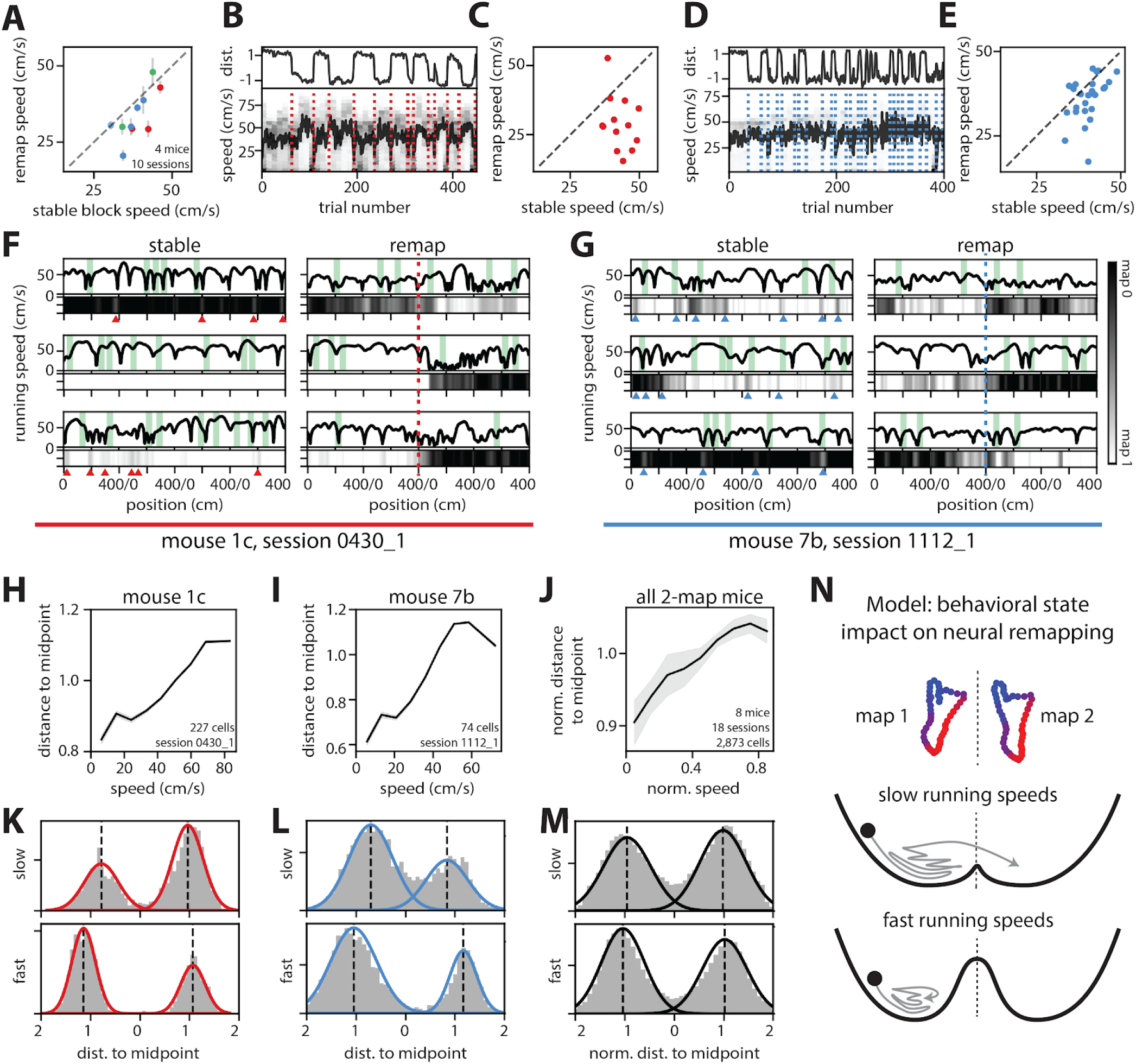
Remapping events and neural variability correlate with slower running speeds. (A) Average running speed in remap trials versus stable blocks (points indicate session average; colors indicate individual mice; grey error bars, SEM; dashed line, unity; n = 10 sessions, 4 mice). (B) (Top) Distance score for population-wide neural activity by trial (as in fig. 2B) for an example cue poor session (n = 227 cells). (Bottom) Running speed by trials (black line, average; gray, density), dotted lines indicate remapping events. (C) as in (A) for the example session shown in (B)(points indicate a remap trial/stable block pair). (D, E) as in (B, C) but for a second example session (cue rich; n = 74 cells). (F) (Top panels) Average running speed by 5 cm position bins for the two trials preceding and following either a remap event (right; dashed line, remap point) or middle of the corresponding stable block (left) for the example session from (B, C)(black trace, running speed; green shading, reward zones). (Bottom panels) Distance to midpoint between k-means clusters by 5 cm position bins (gray, between maps; black, map 0; white, map 1). Arrowheads indicate points of slow running speed where the neural code approached the midpoint. (G) As in (F), but for the example session from (D, E). (H) Average distance to midpoint versus binned running speed for one example session (black line, average; gray shading, SEM). (I) As in (H), but for a second example session. (J) as in (H, I), but for all 2-map sessions; distance to midpoint and speed are normalized within each session (n = 2,873 cells, 18 sessions, 8 mice). (K) Distance to midpoint for 5 cm position bins, split into slow (20th percentile) and fast (80th percentile) average running speeds for an example session (curves, gaussian fit; black dashed lines, means of gaussians; n = 227 cells). (L) As in (K), but for a second example session (n = 74 cells). (M) As in (K, L), but for all 2-map sessions. (N) Schematic model: schematized manifolds (top), neural activity (ball, neural activity; arrow, trajectory in state space), and energy landscape (black line) for slow (middle) and fast (bottom) running speeds shows how running speed might interact with remapping events (dashed line, midpoint between clusters). (See also Fig. S5.)

We next investigated the moment-by-moment correlation between neural variability and running speed by binning neural activity and speed into 5 cm position bins within each trial. For each position bin, we calculated how close the neural activity was to the midpoint between manifolds, where activity is most likely to switch between maps (distance score = 1 in cluster centroid for either map, 0 when equidistant from each manifold)(Methods). As expected, neural activity was closer to the midpoint within remap trials compared to stable blocks (mean distance ± SEM: remap trials = 0.631 ± 0.003, stable blocks = 1.045 ± 0.001; two-sided Wilcoxon rank-sum test, p < 0.0001; n = 80 bins per trial for 226 remap trials, 5,393 stable trials across 18 sessions in 8 mice)(fig. 5f, g bottom). However, we also observed instances where the neural activity approached the midpoint within stable periods (stable bin distance interquartile range: 0.761 to 1.342)(fig. 5f, g arrowheads), indicating that neural variability does not by necessity provoke a remap event. Across all position bins, slower running speeds were correlated with a reduced neural distance to decision boundary (ordinary least squares regression, R^2^ = 0.782, p < 0.0001; n = 10 speed bins per session for 18 sessions in 8 mice)(fig. 5h-j). Further, the medians of the two maps were generally closer in activity space for bins containing the slowest (20th percentile) compared to the fastest (80th percentile) running speeds (11/18 2-map sessions had less separated medians at slow speeds, 7 sessions had more separated medians at slow speeds, p < 0.01)(fig. 5k-m). Altogether, these results suggest that the two neural maps are often less distinguishable at slow running speeds, likely increasing the probability of a remap event. Conceptually, this suggests a model in which remapping involves overcoming an energetic barrier separating two locally stable spatial manifolds. The magnitude of this barrier could change over time and correlate positively with running speed (fig. 5n). In the Supplementary Text, we provide a formal analysis of this hypothesis within the well-established framework of attractor networks (fig. S6a-g).

### Spontaneous remapping persists in double-track sessions

Each of the 2-map sessions was recorded as a single continuous session in just one environment (either cue rich or cue poor). In order to examine remapping in a more dynamic setting, we ran a cohort of mice on alternating blocks of trials in the cue rich and cue poor environments (i.e. “double-track” sessions; n = 13 sessions in 5 mice). Each double-track session consisted of four blocks of 75-100 trials, delineated by 1-2 minutes of darkness (fig. S1). As expected, MEC maintained distinct maps for the distinct environments. Spatial representations from the same environment were qualitatively more similar to one another than to representations of other environment, evidenced by the blocky structure of the network-wide similarity matrices, which matched the blocked trial structure (fig. 6a, b, left; dashed lines indicate breaks between blocks). To assess remapping within each environment, we next divided sessions into trials from either the cue rich (fig. 6a, b, middle) or cue poor (fig. 6a, b, right) track (i.e. “trial blocks”). As trial blocks from the same environment were separated by 75-100 trials, we then re-normalized neural firing rates to correct for small amounts of drift in network-wide spatial representations across these long sessions (see Methods). Similar to single-track sessions, we observed remapping in single neurons (fig. 6c) and across the population (fig. 6a, b middle, d top, i-p), such that each cue rich or poor environment was often represented by several (1-3) stable maps (n = 26 blocks from 13 sessions in 5 mice). We also observed periods of spatial instability (fig. S2). In trial blocks with multiple stable maps, remapping was qualitatively well-captured by a k-means clustering model (fig. 6d, i-p, left bar).

**Figure 6:**
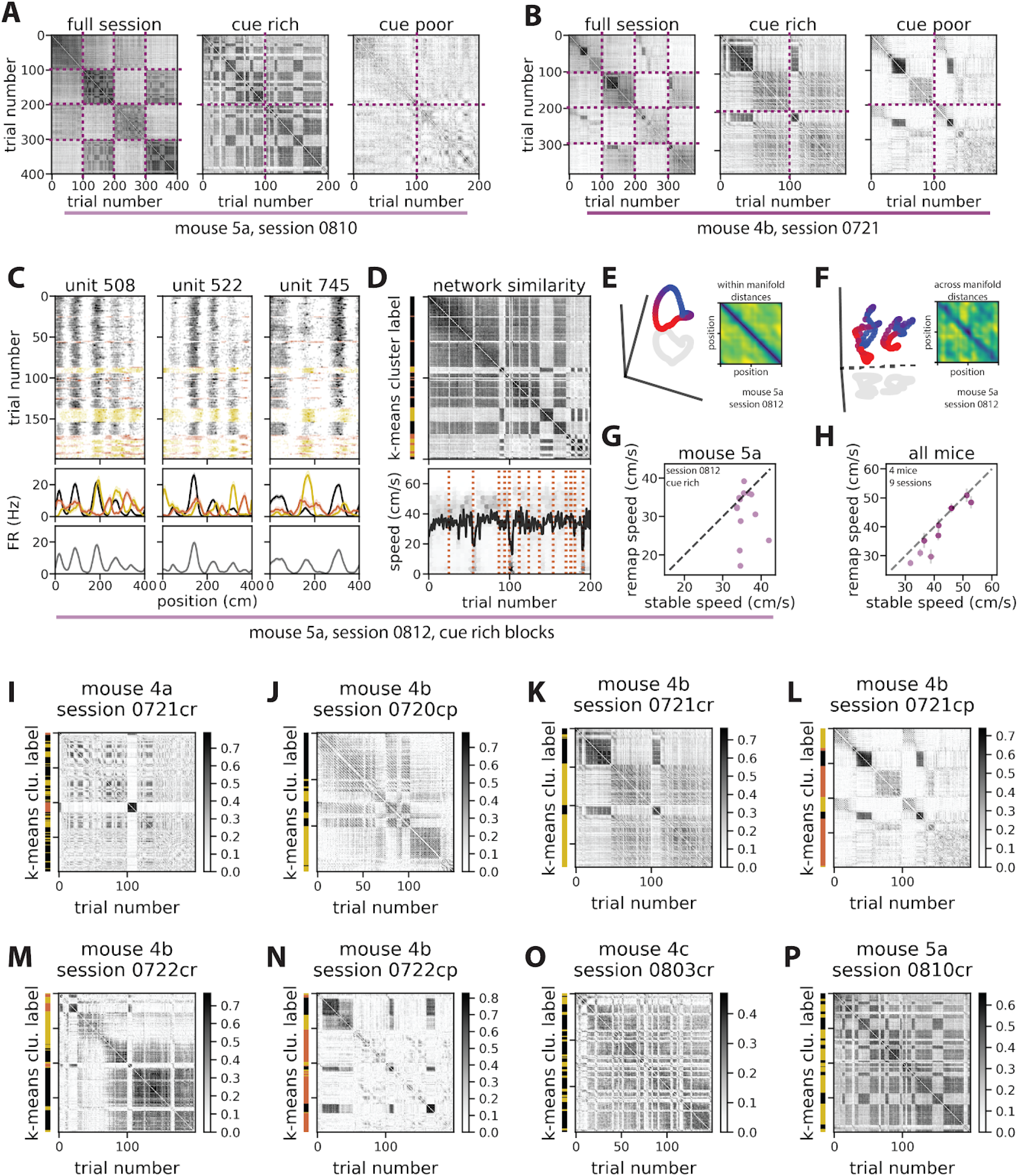
Spontaneous remapping persists in double-track sessions. (A, B) Network-wide similarity matrices for example double-track full sessions (left) and split into cue rich (middle) and cue poor (right) trial blocks (A, n = 68 cells; B, n = 55 cells)(dashed lines indicate breaks between blocks). (C) Single-neuron spiking (top) and tuning curves (middle) for example cells from an example pair of cue rich trial blocks, colored/divided by k-means cluster labels (top), versus averaged over the full session (bottom)(solid line, trial-averaged firing rate; shading, SEM; color scheme denotes map identity and is preserved in D)(compare to Fig. 2F). (D, top) Network-wide spatial similarity for an example pair of cue rich trial blocks (right) and corresponding k-means cluster labels (left). (D, bottom) Running speed by trials (black line, average; gray, density), dotted lines indicate remapping events. (E) PCA projection of a single map (k-means centroid) from an example pair of cue rich trial blocks (n = 130 cells). (Inset) Pairwise distances in neural firing rates across all points (i.e. spatial position bins) in the manifold (color code blue, minimum value; yellow, maximum)(compare to Fig. 4B). (F) PCA projection of two manifolds extracted from an example pair of cue rich trial blocks. (Inset) Across-manifold distances in neural firing rates for every pair of points (color code as in E)(compare to Fig. 4E). (G) Average running speed on remap trials vs. stable blocks for an example session (points, stable block/remap trial pairs; dashed line, unity; n = 13 pairs)(compare to Fig. 5C, E). (H) Average running speed in remap trials versus stable blocks (points indicate session average; colors indicate individual mice; grey error bars, SEM; dashed line, unity; n = 9 sessions in 4 mice). (I-P) As in (D, top), but for additional example 2-map (I, L, M, N) and 3-map (J, K, O, P) double-track sessions (“cr” indicates cue rich blocks from a given session; “cp” indicates cue poor; colorbars indicate trial-by-trial correlation). (See also Fig. S5.)

One feature of the double-track task design is that we recorded from a single population of neurons across repeated visits to the same environment. In this task, the MEC circuit often returned to the same set of maps when returned to the same environment (fig. 6a, b middle, d top, i-p), indicating that the multiple maps were stable over time. Indeed, an example pair of trial blocks, concatenated across repeated visits to the cue rich environment, demonstrates remapping that is virtually indistinguishable from remapping in a continuous single-track cue rich or cue poor session (fig. 6c-g). Single neurons exhibited distinct tuning curves in each map and showed a mix of rate and global remapping (fig. 6c; compare to fig. 2f). Each k-means-identified neural manifold formed a qualitative ring structure in neural activity space (fig. 6e; compare to fig. 4b) and the manifolds from a given session were significantly aligned (versus a randomly rotated null distribution; Methods)(fig. 6f; compare to fig. 4e). Finally, across many double-track sessions, the animal’s average running speed on remap trials was lower compared to its average running speed in the preceding stable block (6/9 sessions; speeds were equal in 3 sessions; mean percent difference in running speed ± SEM: 6.9 ± 2.1%; Wilcoxon two-sided signed-rank test, p = 0.0003; n = 86 remap trial/stable block pairs)(fig. 6d bottom, g, h; compare to fig. 5a-e). Thus our results from continuous single-track sessions appeared to generalize to a task involving both environments.

## Discussion

Here, we report that MEC representations are capable of remapping even in the absence of any changes to sensory cues (Diehl et al., 2017; Fyhn et al., 2007; Marozzi et al., 2015; Solstad et al., 2008) or task demands (Boccara et al., 2019; Butler et al., 2019; Keene et al., 2016). Previous reports have observed that increased running speed is associated with improved spatial tuning in single neurons through sharpening of individual tuning curves (Bant et al., 2020; Hardcastle et al., 2017). Here we show that changes in running speed can also correspond to large shifts in MEC coding—up to three-fold variation in peak firing rate, and 50% reconfiguration in the spatial patterning of neural tuning (fig. 2i-j). Further, we find that remapping events rapidly recruit almost all position-coding neurons across the full anatomical extent of MEC, resulting in synchronous, network-wide remapping events. Finally, these remapping events represent complete transitions between geometrically aligned neural activity manifolds. Together, our results empirically demonstrate that MEC can maintain multiple distinct attractor networks in a single environment, an idea which had previously only been considered theoretically (Sanders et al., 2020).

Consistent with predictions of attractor network models of navigational coding, we find that network activity follows 1D, ring-shaped trajectories through high-dimensional neural state space (Burak and Fiete, 2009; Fuhs and Touretzky, 2006; Guanella et al., 2007; Samsonovich and McNaughton, 1997). Previous attractor models of MEC have demonstrated how, for a given environment, a set of landmark cues can combine with self-motion information to establish a single spatial map and maintain one’s location estimate within that map (Burak and Fiete, 2009; Campbell et al., 2018; Ocko et al., 2018; Skaggs et al., 1995). Theoretical work has also considered how multiple attractors (i.e. internal spatial maps) could be embedded in the same network; however, this possibility is often framed in terms of modeling spatial coding across different external environments (Romani and Tsodyks, 2010; Roudi and Treves, 2008; Samsonovich and McNaughton, 1997; Stringer et al., 2004). Our results build off of these models to suggest that multiple attractors may be visited within the same environment in a rapid and repeating fashion. Further, we find that behavioral factors could influence the probability of transitioning between attractors (Fig. 5n). Importantly, our experiments show that these attractor manifolds are geometrically aligned, suggesting an elegant computational mechanism by which spatial information encoded by MEC could be preserved across remapping events. This geometric alignment could plausibly allow downstream neurons that are insensitive to the remapping dimension to decode position across remapping events, while neurons sensitive to the remapping dimension could decode information unique to each internal context. Future studies will be needed to elucidate how these multiple attractor manifolds are formed and what factors determine the number of distinct maps associated with a particular environment.

Our findings indicate that a change in behavioral state often coincides with network-wide, synchronous remapping in MEC spatial representations. While remapping events often took the form of transitions between two stable spatial maps, we also observed multiple sessions that contained more than two stable maps, as well as sessions where neural coding remapped between landmark-based and landmark-free coding regimes. It is possible that the impoverished sensory experience of our virtual environment promoted these remap events. However, complete remapping of hippocampal spatial representations of a single environment has recently been observed in freely behaving animals, suggesting that similar remapping can occur under more naturalistic settings (Sheintuch et al., 2020). The behavioral-state driven remapping that we observed almost certainly interacts with other factors known to cause MEC remapping, such as changes in the sensory features (Diehl et al., 2017; Fyhn et al., 2007; Marozzi et al., 2015; Solstad et al., 2008) or task demands (Boccara et al., 2019; Butler et al., 2019; Keene et al., 2016) associated with a given environment. It will be of interest for future work to determine the degree to which behavioral state variables versus environmental factors modulate neural variability, with the ultimate goal of predicting which factors will drive remapping across behaviors, time and context. Moreover, detailed consideration and tracking of multiple behavioral state variables will be needed in future work to identify which specific state variables control remapping in the navigation circuitry.

Altogether, we find that MEC has the capacity to remap in a rapid and reversible fashion, which could support a role for this circuit in dividing the unbroken stream of sensory features encountered during navigation into discrete contextual episodes. Further, the current work aligns with a larger body of emerging findings that demonstrate that cortical activity is highly responsive to behavioral state changes (Jennings et al., 2019; Salay et al., 2018; Stringer et al., 2019). Our results suggest that these behavioral state changes may drive rapid, large-scale reconfigurations of internal representations for the external world in higher-order cortex.

## Acknowledgements

We thank A. Diaz for histology assistance and animal care; I. Zucker-Scharff and C. Nnebe for assistance with behavioral training; and A. Attinger, T. Fisher, F.K. Masuda, and J. Wen for discussions and feedback.

## Author Contributions

*Isabel I.C. Low*: Conceptualization, Methodology, Formal analysis, Investigation, Writing - Original draft preparation, Writing - Review & editing, Visualization. *Alex H. Williams*: Conceptualization, Methodology, Formal analysis, Writing - Original draft preparation, Writing - Review & editing, Visualization. *Malcolm G. Campbell*: Conceptualization, Methodology, Writing - Review & editing, Visualization. *Scott W. Linderman*: Conceptualization, Writing - Review & editing, Supervision, Funding. *Lisa M. Giocomo*: Conceptualization, Writing - Original draft preparation, Writing - Review & editing, Supervision, Funding.

## Methods

### Resource Availability

#### Lead Contact

Further information and requests for resources and reagents should be directed to and will be fulfilled by the Lead Contact, Lisa M. Giocomo.

#### Materials Availability

This study did not generate new unique reagents.

#### Data and Code Availability

Data will be made available at https://giocomolab.weebly.com/data.html.

Code will be made available at https://github.com/GiocomoLab.

### Experimental Model and Subject Details

#### Mice

All techniques were approved by the Institutional Animal Care and Use Committee at Stanford University School of Medicine. Recordings were made from 17 C57BL/6 mice aged 4 weeks to 3.5 months at the time of first surgery (15.7–35 g). All mice were female except mice 5a and 5b, which were male. Mice were group housed with same-sex littermates, and in one case with the dam (2a, b, c with 3a), unless separation was required due to water restriction, aggression, or disturbance of prep site. Mice were housed in transparent cages on a 12-h light-dark cycle and experiments were performed during the light phase.

### Method Details

#### Training and handling

Mice were handled at least every 2 days following headbar implantation and given an in-cage running wheel. Starting 1 day after headbar implantation, same-sex mice were placed daily in a large (100×100cm), communal environment with enrichment objects including a running wheel, Lego tower, textured floor tape, and crushed chocolate cheerios for between 15 mins and 1.5 hours. Mice were monitored for aggression and separated as needed. Mice were given free access to water until 3 days after headbar implantation, after which they were water restricted to 1mL of water per day and weighed daily to ensure a body weight of >80% of their starting weight.

After >1 day of water restriction, mice were acclimated to head fixation and trained to drink water from a custom lickport for 10-20 mins over 2 days. Mice were then trained to run on the virtual random forage task (described below) starting with a reward probability of 0.1/cm (essentially one reward per 50 cm), for gradually decreasing reward probability and gradually increasing session length as behavior improved. Mice were trained on the exact track(s) that they were recorded in, either cue poor (litters 1, 2, and 3), cue rich (litters 6 and 7), or both (litters 4 and 5). Mice trained on both tracks were exposed to each track in a counterbalanced fashion, initially alternating tracks over days and ultimately alternating the order of presentation as mice improved sufficiently to run 2 or more sessions per training day. Training continued at least until mice ran >300 trials in 2 hours and demonstrated proficient licking from the lickport in the reward zone; training was sometimes extended in order to stagger recording periods (7 days to 7 weeks; mean ± SEM: 23 ± 3 days; 3 mice never learned the task).

#### *In vivo* survival surgeries

For all surgeries, anesthesia was induced with isoflurane (4%; maintained at 0.5–1.5%) followed by injection of buprenorphrine (0.05–0.1 mg/kg). Animals were injected with baytril (10 mg/kg) and rimadyl (5 mg/kg) immediately following each surgery and for 3 days afterwards. In the first surgery, animals were implanted with a custom-built metal headbar containing two holes for head fixation, as well as with a jewelers’ screw with an attached gold pin, to be used as a ground. The craniotomy sites were exposed and marked during headbar implantation and the surface of the skull was coated in metabond. After completion of training, a second surgery was performed to make bilateral craniotomies (~200μm diameter) at 3.7–3.95mm posterior and 3.3–3.4mm lateral to bregma. A small plastic well was implanted around each craniotomy and affixed with metabond. Craniotomy sites were covered with a drop of sterile saline and with silicone elastomer (Kwik-sil, WPI) in between surgery and recordings.

#### *In vivo* electrophysiological data collection

All recordings were performed at least 16-h after craniotomy surgery, at which point the mouse was head-fixed on the VR recording rig. Craniotomy site was exposed and rinsed with saline—occasionally dura was re-nicked or debris removed using a syringe tip. Recordings were performed using Phase 3B Neuropixels 1.0 silicon probes(Jun et al., 2017) with 384 active recording sites (out of 960 total) along the bottom ~4 mm of a ~10 mm shank (70 μm wide shank diameter, 24 μm thick, 25 μm electrode spacing), and reference and ground shorted together. The probe was positioned over the craniotomy site at 8–14° from vertical and targeted to ~50–300 μm anterior of the transverse sinus using a micromanipulator. On consecutive recording days, probes were targeted medial or lateral of previous recording sites as permitted by the craniotomy. The reference electrode was then connected to a gold ground pin implanted in the skull. The probe was advanced slowly (~10 μm/s) into the brain until it encountered resistance or until activity quieted on channels near the probe tip, then retracted 100–500μm and allowed to sit for at least 30 mins prior to recording. While the probe was implanted, the craniotomy site was covered with sterile saline and silicone oil. Signals were sampled at 30 kHz with gain = 200 (7.63 μV/bit at 10 bit resolution) in the action potential band, digitized with a CMOS amplifier and multiplexer built into the electrode array, then written to disk using SpikeGLX software.

#### Virtual reality environment

The VR recording set-up was nearly identical to that in Campbell et al (Campbell et al., 2018). Head-fixed mice ran on a 15.2-cm-diameter foam roller (ethylene vinyl acetate) constrained to rotate about one axis. The cylinder’s rotation was measured by a high-resolution quadrature encoder (Yumo, 1024 P/R) and processed by a microcontroller (Arduino UNO). The virtual environment was generated using commercial software (Unity 3D) and updated according to the motion signal. Virtual reality position traces were synchronized to recording traces on each frame of the virtual scene. The virtual scene was displayed on three 24-inch monitors surrounding the mouse. The gain of the linear transformation from ball rotation to translation along the virtual track was calibrated so that the virtual track was 4 m long. At the end of the track, the mouse was teleported seamlessly back to the start to begin the next trial, such that the track was seemingly infinite (all visual cues were repeated and visible into the distance as the mouse approached the track end).

Cue rich tracks consisted of 5 towers of different heights, widths, and patterns (black and white, neutral luminance), placed at 80 cm intervals starting at 0 cm (see schematic, fig. 1c, top) and a black and white checkerboard on the floor for optic flow. Cue poor tracks contained 2 towers of different patterns placed at 0 and 200 cm (see schematic, fig. 1c, bottom) and a white to black horizontal sinusoidal pattern on the floor. Both tracks had uniform gray walls and sky beyond the towers. For mice that experienced a single track, recording sessions consisted of 57–450 trials (mean ± SEM: 328 ± 14 trials). For mice that experienced both tracks, each track was presented in a block of 75–100 trials with ~1 min of darkness in between tracks (fig. S1g). Blocks alternated between cue rich and poor—each track was presented twice (barring rare cases when the mouse failed to complete the session) and which track was presented first was alternated on each day.

#### Random foraging task

In both cue rich and cue poor tracks, visually marked reward zones appeared at a probability of 0.01–0.001 per cm, titrated to mouse performance, within the middle 300 cm of the track and at least 50 cm apart. Reward zones were 50 cm long and track-width, were patterned with a black and white diamond checkerboard, and hovered slightly above the floor. Upon entering the reward zone, animals could request water by licking and breaking an infrared beam at the mouth of the lickport; if not requested, water was dispensed automatically at the center of the zone. For mice 1c, 4a, and 4b for some recording sessions there were between 1-5 probe trials every 10 trials in which water was only dispensed if requested in the reward zone (no automatic dispensation). Upon water dispensation (or next trial start for missed probe trials), the current reward zone disappeared and the next zone became visible. Water rewards (~1.5 μL) were delivered using a solenoid (Cole-Parmer) triggered from the virtual environment software, generating an audible click with water delivery. Licks were recorded as new breaks in the lickport infrared beam and were processed by a microcontroller (Arduino UNO).

#### Histology and probe localization

Before each implantation, probes were dipped in fixable lipophilic dye (1mM DiI, DiO, DiD, Thermo Fisher) 10 times at 10 second intervals. Within 7 days of the first probe insertion, mice were killed with an overdose of pentobarbital and transcardially perfused with phosphate-buffered saline (PBS) followed by 4% paraformaldehyde. Brains were extracted and stored in 4% paraformaldehyde for at least 24 h before transfer to 30% sucrose in PBS. Brains were then rapidly frozen, cut into 30-μm sagittal sections with a cryostat, mounted and stained with cresyl violet. Histological sections were examined and the location of the probe tip and entry into the dorsal MEC for each recording were determined based on the reference Allen Brain Atlas (Allen Institute for Brain Science, 2004)(Fig. S1). The location of each recording site along the line delineated by the probe tip and entry point was then determined based on each site’s distance from the probe tip. Only cells within MEC, again based on the reference Allen Brain Atlas (Allen Institute for Brain Science, 2004), were included for analysis (Fig. S1). In all cases, “depth” reported is the ventral distance from the location of the dorsal boundary of MEC in the medial section where the probe enters MEC.

#### Offline spike sorting

Electrophysiological recordings were common-average referenced to the median across channels and high-pass filtered above 150 Hz. Automatic spike sorting was then performed using Kilosort2, a high-throughput spike sorting algorithm that identifies clusters in neural data and is designed to track small amounts of neural drift over time (open source software by Marius Pachitariu, Nick Steinmetz, and Jennifer Colonell, https://github.com/MouseLand/Kilosort2)(see also Kilosort1 (Pachitariu et al., 2016)). After automatic spike-sorting, all clusters with peak-to-peak amplitude over noise ratio < 3 (with noise defined as the standard deviation of voltage traces in a 10ms window preceding detected spike times), total number of spikes < 100, and repeated refractory period violations (0-1 ms autocorrelegram bin > 20% of maximum autocorrelation) were excluded. All remaining clusters were manually examined and labeled as “good” (i.e. stable and likely belonging to a single, well-isolated neural unit), “MUA” (i.e. likely to represent multi-unit activity), or “noise.” Only well-isolated “good” units from within MEC (barring fig. S1h, i, which were non-MEC units) with greater than 400 spikes were included for analysis in this paper. Sessions with fewer than 10 cells meeting these criteria were excluded.

#### Behavioral data preprocessing

On each frame of the virtual reality scene, the virtual position and time stamps were recorded and a synchronizing TTL pulse was sent from an Arduino UNO to the electrophysiological recording equipment. These pulses were recorded in SpikeGLX using an auxiliary National Instruments data acquisition card (NI PXIe-6341 with NI BNC-2110). The location of each reward zone, time of each lick, and time of each reward dispensation were also recorded. Thus all time stamps and behavioral factors were synchronized to the neurophysiological data. Time stamps were adjusted to start at 0 and all behavioral data was interpolated to convert the variable VR frame rate to a constant frame rate of 50Hz. As the track was effectively circular and 400 cm long, recorded positions less than 0 or greater than 400 cm were converted to the appropriate position on the circular track (eg. a recorded position of 404 cm would be converted to 4 cm and a recorded position of −4 cm would be converted to 396 cm). Trial transitions were identified as timepoints where the difference in position across time bins was less than −100 cm (i.e. a transition from ~400 cm to ~0 cm) and a trial number was accordingly assigned to each timepoint.

Running speed for each timepoint was computed by calculating the difference in position between that timepoint and the previous, divided by the framerate (speed at the first timepoint was assigned to be equal to that at the second timepoint). Speeds greater than 150 cm/s or less than −5 cm/s were removed. Speed was then interpolated to fill removed timepoints and smoothed with a gaussian filter (standard deviation 0.2 time bins). For all analyses except lick and reward zone analyses (fig. 1d, e, g, h), stationary time bins (speed < 2 cm/s) were excluded.

#### Population analysis and clustering model

The 1D track was divided into 5 cm position bins (total of 80 bins). On each traversal of the track, the empirical firing rate of each neuron—i.e., number of spikes divided by time elapsed—was computed for every position bin. We then smoothed the firing rate traces with a Gaussian filter (standard deviation 5 cm). For each session this resulted in a 3-dimensional array of raw firing rates, with dimensions corresponding to trials, positions, and neurons.

Since these raw firing rates varied widely across neurons, we rescaled them so that the peak firing rate was commensurate across cells. Similar normalization steps or variance-stabilizing transformations have been used in previous population analyses of neural data (Churchland et al., 2012; Williams et al., 2018; Yu et al., 2009), to prevent neurons with high firing rates from washing out low firing rate neurons. Here, we normalized firing rates by first clipping the maximum firing rate of each neuron at its 90th percentile (to exclude large outliers), and then re-scaling each neuron’s firing rate to range between zero and one. That is, if 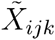 denotes the clipped firing rate on trial *i*, position bin *j*, and neuron *k*, then the normalized firing rate was computed as:

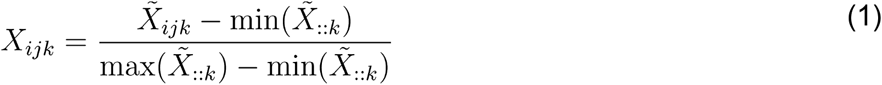

The max(·) and min(·) operations (as well as the 90th percentile clipping operation) are applied on a neuron-by-neuron basis, pooling observations across all trials and timebins. For two-track sessions (which contained 2 blocks of cue rich trials and 2 blocks of cue poor trials, in alternating order) we added an additional per-neuron correction factor to account for drift in firing rates: the mean normalized firing rate for each neuron (across all trials and position bins) was subtracted within each block of trials, and the result was renormalized to values between zero and one, as above.

On each trial, MEC’s representation of position is summarized by a matrix, denoted *X*_*i*::_ for trial *i*, with rows and columns respectively corresponding to position bins and neurons. A simple measure of similarity between two trials, indexed by *i* and *i′*, is given by the Pearson correlation between the vectors vec(*X*_*i*::_) and vec(*X*_*i′*::_). Network-wide trial-by-trial similarity matrices (as in fig. 1, 2, S1, S2, S3, and S5) were found by computing this correlation across all pairs of trials.

Let *X*_*ijk*_ denote the *I* × *J* × *K* array, or tensor, of normalized firing rates defined in equation (1). As before the index variables *i*, *j*, and *k*, respectively represent trials, position bins, and neurons. Now consider the following low-rank matrix factorization model (Singh and Gordon, 2008; Udell et al., 2016) of these data:

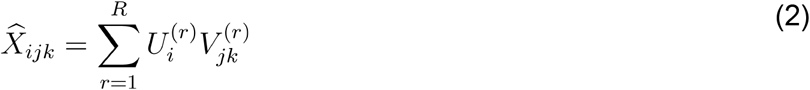

where *R* < min({*I*, *J*, *K*}) denotes the number of model components, or the model rank. Equation (2) represents an approximate factorization of the *I* × *JK* matricization or tensor unfolding of the data array (Kolda and Bader, 2009; Seely et al., 2016)). We will see that k-means clustering arises as a special case of this model, and in this special case represents the number of clusters (i.e. the number of spatial maps).

The parameters 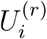 and 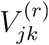 are optimized according to a least squares criterion, i.e.:

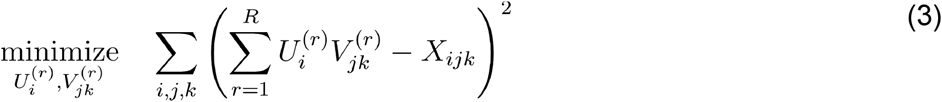

It is well-known that a rank-R truncated singular value decomposition (SVD) provides a solution to this optimization problem (Eckart and Young, 1936). Further, the solution provided by truncated SVD is closely related to Principal Components Analysis (PCA)—indeed, these two methods are identical for the case of mean-centered data (see, e.g., (Shlens, 2005)). Since the normalized firing rate array in *X*_*ijk*_ equation (3) is not mean-centered, we refer to this initial model as “uncentered PCA.” We use the uncentered coefficient-of-determination (uncentered R^2^) as a normalized measure of model performance associated with equation (3).

The k-means clustering model incorporates an additional constraint into the uncentered PCA model. Specifically, k-means seeks to minimize equation (3),

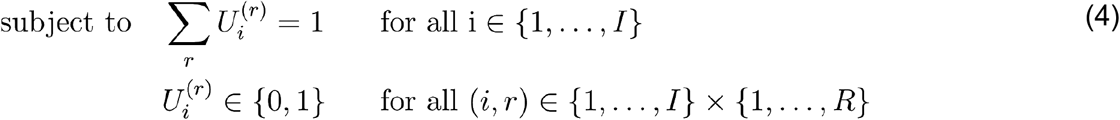

Thus, if we view 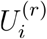 as the elements of an *R* × *I* matrix, the rows of this matrix are constrained to be standard Cartesian basis vectors of *R*-dimensional Euclidean space. Each of these vectors specifies the cluster assignment label for every trial (see fig. 2a for a schematic illustration for the *R* = 2 case). Further, we can interpret 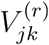 as elements of an *R* × *J* × *K* array. For a fixed cluster index *r*, the remaining elements form a *J* × *K* matrix, called a “slice” of the original array (Kolda and Bader, 2009). Each slice corresponds to a cluster centroid, which we may interpret as a spatial map—the columns are *J* dimensional vectors holding the spatial tuning curves for every neuron, so different slices correspond to different sets of spatial tuning curves for each neuron.

This connection between k-means clustering and other matrix factorization models is well-known and expanded upon in detail by (Singh and Gordon, 2008; Udell et al., 2016)). We exploit this connection to assess the k-means model, which is more constrained than uncentered PCA (i.e. truncated SVD) in that *U*_*i*_ is constrained to be a one-hot vector as opposed to an arbitrary length-R vector. Intuitively, this allows us to interpret each trial as belonging to one of *R* types, as opposed to a linear combination of them. The fact that the more restrictive k-means model performs as well as uncentered PCA gives credence to the multiple-map interpretation. To compare these two models we used a randomized cross-validation procedure in which 10% of the data, representing the validation set, were censored in a speckled holdout pattern (Williams et al., 2018; Wold, 1978). Ten randomized replicates were performed for all models. For the case of *R* = 2 components, we observed similar performance (measured by the uncentered R^2^ averaged over validation sets) between uncentered PCA and k-means for all sessions (see fig. 2d).

Further, we compared the test performance of k-means on “shuffled” datasets (see fig. 2c, fig. 2e). Firing rates from a behavioral session were shuffled by applying a random rotation (i.e., an orthogonal linear transformation) to *X*_*ijk*_ across trials. That is, we sample a random rotation matrix *Q*_*ii′*_ and define

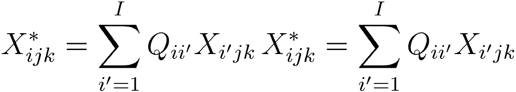

as the new shuffled dataset, which is substituted into the objective function defined in equation (3). This form of shuffling preserves many features of the data, including the overall norm of the data and correlations between neurons and position bins. However, it destroys the sparsity pattern on 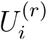 which is imposed by the k-means model. This procedure is similar in spirit to methods proposed by Elsayed & Cunningham (Elsayed and Cunningham, 2017).

Sessions that were well-approximated by the k-means model *R* = 2 with clusters were classified as “two-map” sessions. We required that the performance gap (measured by the uncentered R^2^ averaged over validation sets) between k-means and uncentered PCA and be less than 70% relative to the shuffle control. Further, we required an uncentered R^2^ of at least 0.63 for the k-means model with *R* = 2 maps. Sessions not meeting these criteria sometimes displayed more than two maps (see Fig S6), long periods of unstable coding (see Fig S2), or little to no remapping at all. For all k-means analyses, we ran the clustering model at least 100 times on all neural data from each session to account for model variability, keeping the model with the best fit to the data.

#### Manifold alignment analysis

We used standard Procrustes analysis methods (Gower et al., 2004) to assess the degree to which the two ring manifolds, representing spatial maps, were aligned in neural firing rate space. Recall that the k-means centroids provide an estimate of each spatial map—in this case, we restrict our focus to *R* = 2 maps, so the two maps are given by 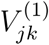 and 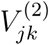. Geometrically, these maps are represented as points *J* embedded in a *K*-dimensional space (recall that *J* denotes the number of position bins and *K* is the number of simultaneously recorded neurons). Procrustes Analysis begins by centering each of these manifolds at the origin and rescaling them to have unit norm. Let 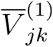 and 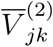 denote the maps after these preprocessing steps have been applied, i.e.,

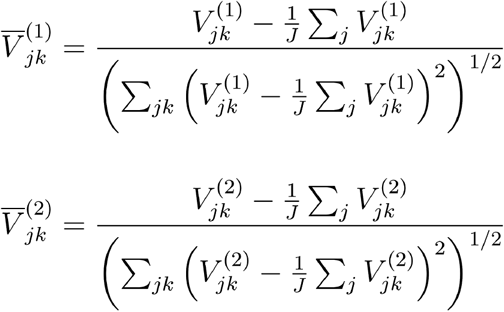

Since position bin *j* in map 1 and position bin *j* in map 2 correspond to the same location on the track, we consider the root-mean-squared-error (RMSE) between these centered and rescaled maps as the empirically “observed” alignment score (reported in fig. 4j):

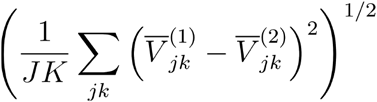

The central step of Procrustes analysis is to find the optimal rotation matrix that aligns these two point clouds. That is, we wish to find the matrix 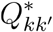 that solves the following optimization problem: 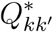 that solves the following optimization problem:

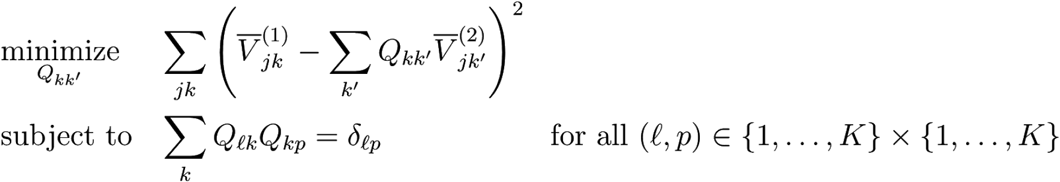

This optimization problem admits a closed form solution for 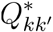 in terms of the singular value decomposition of the *K* × *K* matrix 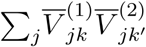 (Schönemann, 1966). See Gower & Dijksterhuis (Gower et al., 2004) for further details and background. In fig. 4j, we report the RMSE after applying the optimal rotation (“aligned”) as well as a random rotation matrix (“shuffled”).

#### Manifold entanglement

We quantified the entanglement of a manifold as the maximum ratio of intrinsic distance (i.e. distance along the manifold) to extrinsic distance (i.e. Euclidean distance in *K*-dimensional space) between any two points on the manifold. Concretely, the extrinsic distance between two points corresponding to position bins and was computed as:

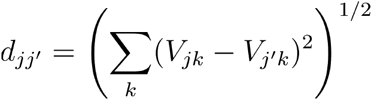

The intrinsic distance, *D*_*jj′*_, was the sum of extrinsic distances along a path from and *j* and *j′* (see diagram in fig. 4d). Depending on whether one travels clockwise or counterclockwise along the ring, there are two paths connecting any pair of points—the intrinsic distance is given by whichever path is shorter. The triangle inequality implies that *D*_*jj′*_≥*d*_*jj′*_ for every pair of points along the manifold. A raw measure of entanglement is given by max[*D*_*jj′*_/*d*_*jj′*_]. We normalized the entanglement scores to values between zero and one by computing:

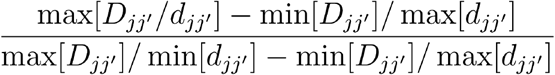

All max and min operations are understood to operate over all pairs of position bins, *j* and *j′*.

#### Distance to cluster calculations

After fitting the k-means model and obtaining two centroids, 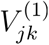 and 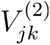, we can quantify how close network activity is to each centroid on a trial-by-trial or neuron-by-neuron basis. In each case we project the activity onto a one-dimensional space where −1 corresponds to one centroid and +1 corresponds to the other centroid. That is, for each trial *i*, we compute

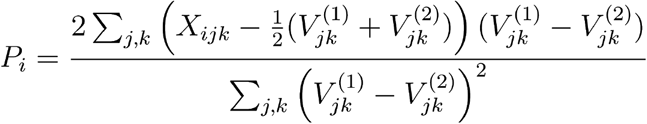

Note that *P*_*i*_ = 1 when 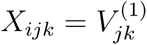 and *P*_*i*_ = −1 when 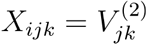. Further, if the network activity on trial *i* is at the midpoint, then 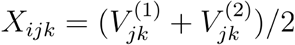 and *P*_*i*_ = 0.

We can compute an analogous statistic for each combination of trial *i* and position bin *j*:

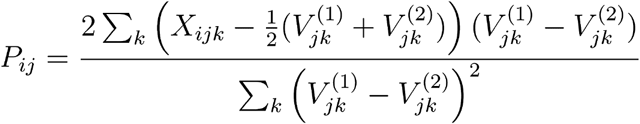

Likewise, we can compute for each combination of trial *i* and neuron *k*:

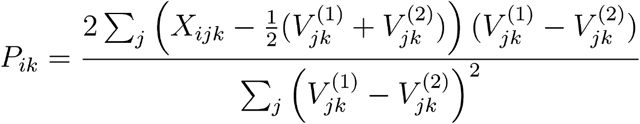

Note that *P*_*i*_, *P*_*ij*_, and *P*_*ik*_ refer to three different quantities, and are only distinguished on the basis of their indices. This concise, somewhat informal, notation is common in tensor algebra, but is restricted to the present section to prevent potential confusion.

In Figure 2, we use *P*_*ik*_ to identify neurons that consistently remap. Let *c*_*i*_ denote the cluster label of each trial such that *c*_*i*_ = 1 if trial *i* is in map 1 and *c*_*i*_ = −1 if trial *i* is in map 2. Then log(1 + exp(−*c*_*i*_*P*_*ij*_) provides a measure of neuron *k*’s distance from the cluster centroid on trial *i* (specifically, it corresponds to a logistic loss function in the context of classification models). Averaging this distance over trials summarizes the remapping strength—intuitively, an average distance close to zero implies that the neuron “agrees with” the rest of the population on each trial, while a large average distance implies that the neuron is inconsistent (e.g., because the neuron exhibits high levels of noise combined with little to no changes in spatial tuning across maps). We classified neurons as “consistent remappers” when the average distance was less than 1. In fig. 5, we used *P*_*ij*_ to assess the relationship between running speed and the distance of neural coding to the midpoint between clusters. In fig. 5h-i, we plotted |*P*_*ij*_|, i.e. the distance to midpoint, in 10 running speed bins—the first 9 bins were evenly spaced between 0 cm/s and 20 cm/s below the maximum speed; the last bin included all top speeds above this final threshold (this was done to account for rare bursts of high speeds). Similarly, in fig. 5j we plotted |*P*_*ij*_|, normalized within each session, in 10 running speed bins, also normalized within each session. In fig. 5k-m, we use histograms to visualize *P*_*ij*_ for all trials and position bins. To account for arbitrary map assignment, we randomly flipped the sign of *P*_*ij*_ for each session in fig. 5m. Likewise, the white-to-black heatmaps in fig. 5f-g visualize *P*_*ij*_ for a subset of trials.

#### Position decoding analysis

We fit linear models to predict the animal’s position from the spiking activity of all MEC neurons, and call the optimized model a “decoder” following common terminology and practice (Kriegeskorte and Douglas, 2019) (fig. 3). Let *y*_*t*_ ∈ [0, 2π) denote the position on the circular track at time *t*, and let *s*_*nt*_ denote the number of spikes fired by neuron *n* at time *t* after smoothing with a Gaussian filter (standard deviation = 200 ms). Due to the nature of the VR environment, *y*_*t*_ is a circular variable—i.e., it should be interpreted as an angle on the unit circle. In the statistics literature, a regression that predicts a circular variable from linear covariates is known as a *circular-linear regression* model. Several approaches to circular-linear regression have been developed (Fisher and Lee, 1992; Pewsey and García-Portugués, 2020; Sikaroudi and Park, 2019). Here, we used a *spherically projected multivariate linear model* (Presnell et al., 1998). Two regression coefficients, 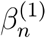 and 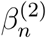, are optimized for each neuron using the expectation maximization routine described by Presnell et al. (Presnell et al., 1998). After fitting the model to a set of training data, the model estimate for a given set of inputs is given by

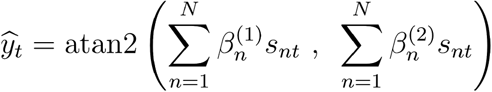

where atan2 corresponds to the “2-argmuent arctangent” function. The “model score” referenced in Figure 2 is the average of 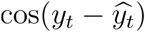 over time bins in the testing set. Thus, a decoder which randomly guesses angles over the unit circle would have an expected score of zero, while a perfect decoder would have a score of one. Note that training data was downsampled to match spike number, position bins, running speed, and number of observations across training sets (map 0, map 1, and both maps) for each session. In fig. 3d, each point represents the test set (10% of the subsampled data) model score for a single training set (the remaining 90% of the subsampled data), while in fig. 3e-g each point represents the average model score across all 10 possible test sets.

#### Spike waveform analysis

Waveforms were extracted from the Kilosort2 data output using Daniel J. O’Shea’s neuropixel-utils library (open source code from GitHub: https://djoshea.github.io/neuropixel-utils/). To compute the spike-by-spike similarity matrices in fig. S4 (C, H, K, O), we extracted the waveforms for 1000 spikes from random points in the session (or all spikes, for n total spikes < 1000) on the 20 best channels. We then concatenated these waveforms *I* × *JK* into an matrix, where *I* = 1000 (spikes), *J* = 20 (channels), and *K* = 82 (samples). We then computed the Pearson correlation between all *I*(*I* + 1)/2 unique pairs of rows and then visualized these results in spike-by-spike similarity matrices (this overall procedure is analogous to how trial-by-trial similarity was computed in fig. 1, 2, S1, S2, S3, and S5). The amplitude plots in fig. S4 (D, I, N, R) represent the peak amplitude for all spikes, as calculated by Kilosort2.

In order to explicitly compare waveforms across remap events and across the session (as in fig. S4 B, G, M, Q), we identified stable blocks of 10 or more trials abutting k-means-identified remap events near the beginning, middle, and end of the session (fig. S4, A, F, L, P). We then extracted the waveforms for 100 spikes randomly chosen from the 10 trials in the middle of each stable block (for cells with fewer than 100 spikes in any pre- or post-remap stable block, we extracted an equal number of spikes from each block, equal to the minimum number of spikes across the two blocks). We then computed the average waveform within each block (e.g. fig. S4, B, G).

#### Tetrode recordings and analysis

The tetrode data included in figure S4 were collected for a previous publication (n = 296 cells from 112 sessions in 19 mice)(Campbell et al., 2018). Because VR gain manipulations as performed for that study can induce remapping of spatial representations, all data examined for this figure were from “baseline trials” in which no VR manipulation occurred. However, because these mice experienced frequent gain manipulations, we compared these tetrode data to Neuropixels data collected from mice who also experienced gain manipulations, to account for potential lasting effects of the manipulations (n = 3,075 cells from 89 sessions in 20 mice)(Campbell et al., 2020). For each cell (Neuropixels or tetrodes), we examined a single block of 20 baseline trials in which no VR manipulation occurred. For each cell, we computed firing rate maps in single trials and computed a cross correlation matrix over the 20 trial block, taking the peak cross correlation over lags from −20 cm to +20 cm to allow for small shifts. We focused our analysis on cells that were "spatially stable" within the first 6 trials (defined as having mean trial-trial peak cross correlation > 0.5 in the first 6 trials), and asked how the rate maps changed in the following 14 trials. To quantify this, we computed the peak cross correlation between each of these 14 trials and each of the 6 "baseline" trials, and averaged over baseline trials. When the pattern remaps, this cross correlation should be low; when it is stable, it should be high. We performed a statistical comparison of the distribution of cross correlations for cells recorded with tetrodes to the distribution for cells recorded with Neuropixels probes. Note that the tetrode recordings were performed on a VR track with more salient visual landmarks than that used for Neuropixels recordings, including a clearly delineated trial structure with teleportations between each trial.

#### Spatial information calculations

Following the procedures in Skaggs et al. (1996)(Skaggs et al., 1996) we calculated spatial information content in bits per second over 2 cm position bins for each neuron as follows:

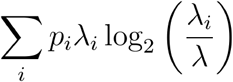

Where *p*_*i*_ is the probability that the animal is in position bin *i* (occupancy of position bin *i* / total session time), λ_*i*_ is the average firing rate of the neuron in position bin *i*, and λ is the overall average firing rate for the neuron. Firing rates were computed empirically (number of spikes / occupancy). For fig. 2, spatial information was calculated separately for each map by first separating trials by their k-means-defined cluster label. Cells were defined as significantly spatial if their spatial information score was > 95^th^ percentile of a null distribution comprising 1,000 shuffle controls. Shuffles were computed separately for each map in each session, and were implemented by shifting all spikes for all cells by a random time interval, up to a maximum of 10 seconds, to disrupt the spike/position relationship without changing inter-spike intervals or correlations across cells. “Spatial cells” in fig. 2 and as referred to throughout the text, were significantly spatial in at least one map.

#### Rate remapping vs global remapping

In order to identify the extent to which each cell remapped across trials, we first divided trials by their 2-factor k-means cluster label. We then computed the trial-averaged firing rate in 2 cm position bins for each map, smoothing with a 1D Gaussian filter (standard deviation 2 cm). We quantified the degree of rate remapping in each neuron by the percent change in the peak firing rate (i.e. largest firing rate in any spatial bin) across the two maps. As a measure of global remapping, we calculated an alignment score between the normalized firing rate vectors in activity space. To do so, we computed the cosine similarity (vector dot product after normalization) between the spatial profiles of the within-map averaged firing rates. A cosine similarity of 1 would indicate an identical spatial firing pattern and a score of 0 would indicate orthogonal spatial representations.

#### Spatial autocorrelation analysis

In fig. S2, we analyzed periods of unstable spatial coding in MEC (i.e. periods where spatial tuning curves were not entrained to landmark locations). We first divided trials into spatial and non-spatial using a threshold value of 0.4 average spatial correlation to the 4 nearest trials (using the trial-by-trial spatial similarity matrix described in “Population analysis and clustering model”). For each trial, we computed the firing rate in 2 cm position bins and concatenated trials of each type to obtain a continuous vector of firing rate by absolute distance traveled for each condition. We then computed the spatial autocorrelation of this signal, up to a maximum distance of 1600 cm, normalized such that the autocorrelation at 0 cm = 1. We identified peaks in this signal with a minimum prominence of 0.05 and compared each peak to a null distribution to determine whether it was higher than could be expected by chance. The null model was given by a first-order autoregressive process, i.e. an AR(1) model, with no drift term. The decay parameter of the model was chosen to be 0.55 to roughly match the autocorrelation of cells with small spatial fields. The autocorrelation function of the AR(1) null model admits a closed-form expression, as covered in standard references on time series analysis(Chatfield, 1984). To evaluate whether any peak in the empirical autocorrelation function was significant with respect to this null distribution, we computed Bonferroni-corrected 95% confidence intervals around the observed autocorrelation function. We classified a given neuron as having periodic firing fields if it had more than one significant peak in the unstable regime.

### Quantification and Statistical Analysis

#### Statistics

All data were analyzed in Python, using the scipy stats library to compute statistics, except for data in fig. S4u, which were analyzed in MATLAB. Unless otherwise noted, all tests are two-sided, correlation coefficients represent Pearson’s correlation, and values are presented as mean ± standard error of the mean (SEM). Non-parametric tests were used to assess significance, specifically Wilcoxon signed-rank tests for paired data, Wilcoxon rank-sum tests for unpaired data, and Kruskal-Wallis H-tests for comparisons of > 2 values. Data collection and analysis were not performed blind to the conditions of the experiments. No statistical methods were used to predetermine sample sizes, but our sample sizes are consistent with previous similar studies.

## Supplementary Information

**Fig. S1:**
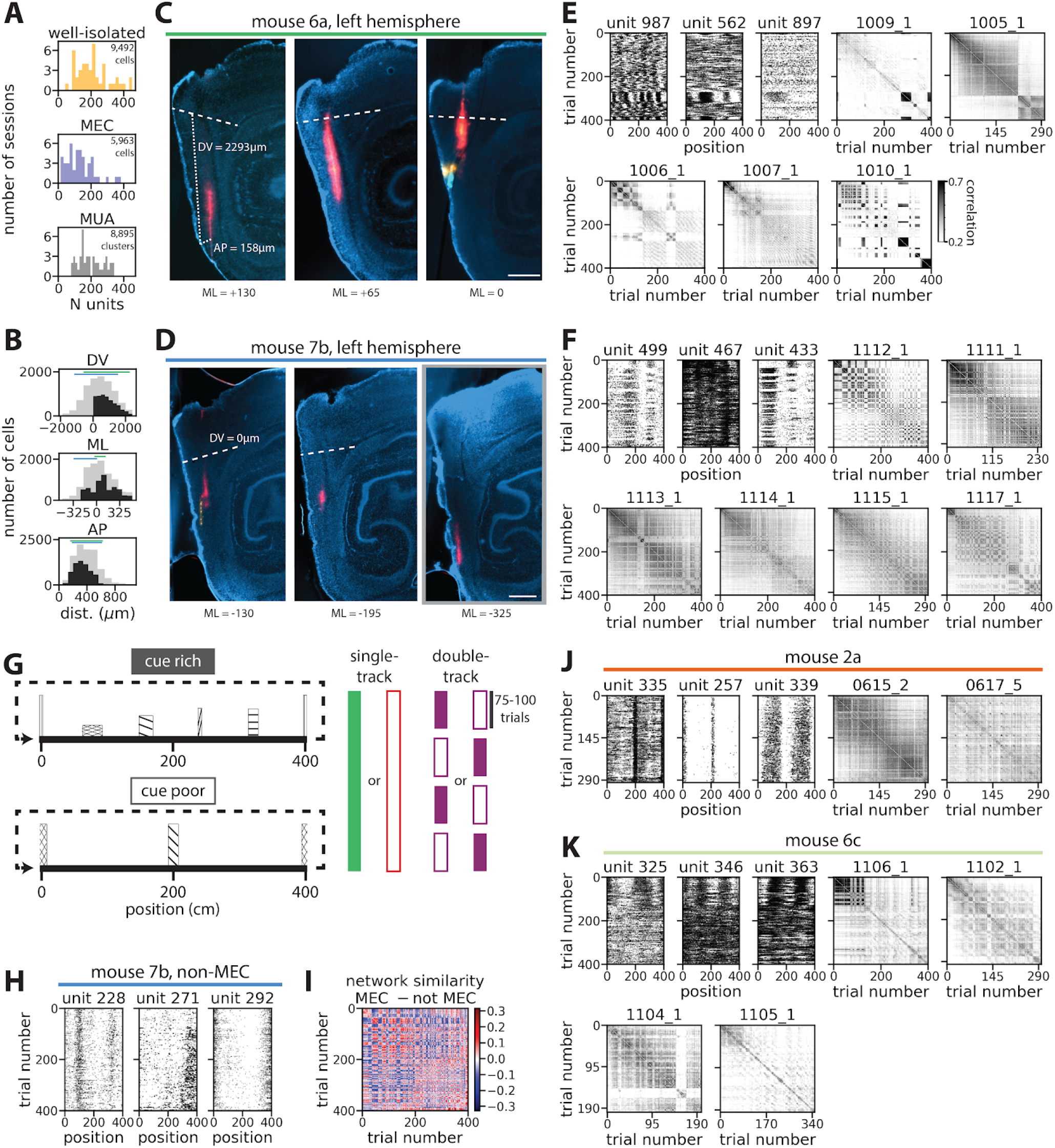
Unit identification, histology and examples of remapping in individual mice and sessions. (A) Number of kilosort-identified clusters from each session labeled as well-isolated single units (top, total cell n = 9,307), the subset of those units within MEC (middle, total cell n = 5,844), and the number of clusters labeled as multi-unit activity (“MUA”, bottom, total cluster n = 8741) for all sessions with >10 MEC units (total cluster/unit n = 18,048 from 44 sessions in 17 mice). (B) Locations for all recorded units relative to anatomical boundaries (black, MEC units; gray, all well-isolated units; bars, probe location for example mice shown in panels C and D). Note that units from the same dorsal-ventral (DV), medial-lateral (ML), or anterior-posterior (AP) coordinates can be classified as either within or outside of MEC. For example, see (D, far right panel, red dye) for probe placement with acceptable DV and AP coordinates, but unacceptable ML coordinates (−325μm ML is too far medial to be within MEC). (C) Example histology from mouse 6a, left hemisphere showing probe placement for session 1009_1 (red)(bottom text, medial (−) or lateral (+) distance from center of MEC (ML = 0μm); dashed lines, dorsal MEC boundary (DV = 0μm); scale bar = 500μm). AP distance is calculated as the perpendicular distance from probe tip (or probe crossing of MEC boundary) to the back of the brain (AP = 0μm); DV location is determined as distance from that point at the back of the brain to the MEC dorsal boundary, traveling parallel to the probe track (C left, dotted lines). (D) As in (C), but for mouse 7b, left hemisphere (red, session 1112_1). (E) Trial-by-trial similarity matrices for all sessions from mouse 6a with >10 MEC units; raster plots (top left) show example units from session 1009_1. (F) As in (E) but for mouse 7b, raster plots (top left) show example units from session 1112_1. (G) Schematic of cue rich and poor tracks (left) and task structure for single-track (middle) and double-track (right) sessions. (H) Three example cells from the example session in (D, F) recorded outside MEC illustrate that non-MEC cells tended not to show remapping. (I) In order to compare network-wide remapping for MEC cells to non-MEC cells recorded in the same session, we subtracted the trial-by-trial similarity matrix for non-MEC cells from the trial-by-trial similarity matrix for MEC cells. Within map similarity was overall higher and across map similarity was overall lower for MEC cells compared to non-MEC cells, indicating stronger remapping in this population (colorbar indicates the difference in similarity between MEC and non-MEC trial-by-trial similarity). (J, K) Rasters and similarity plots from additional example mice and sessions show a variety of remapping patterns.

**Fig. S2:**
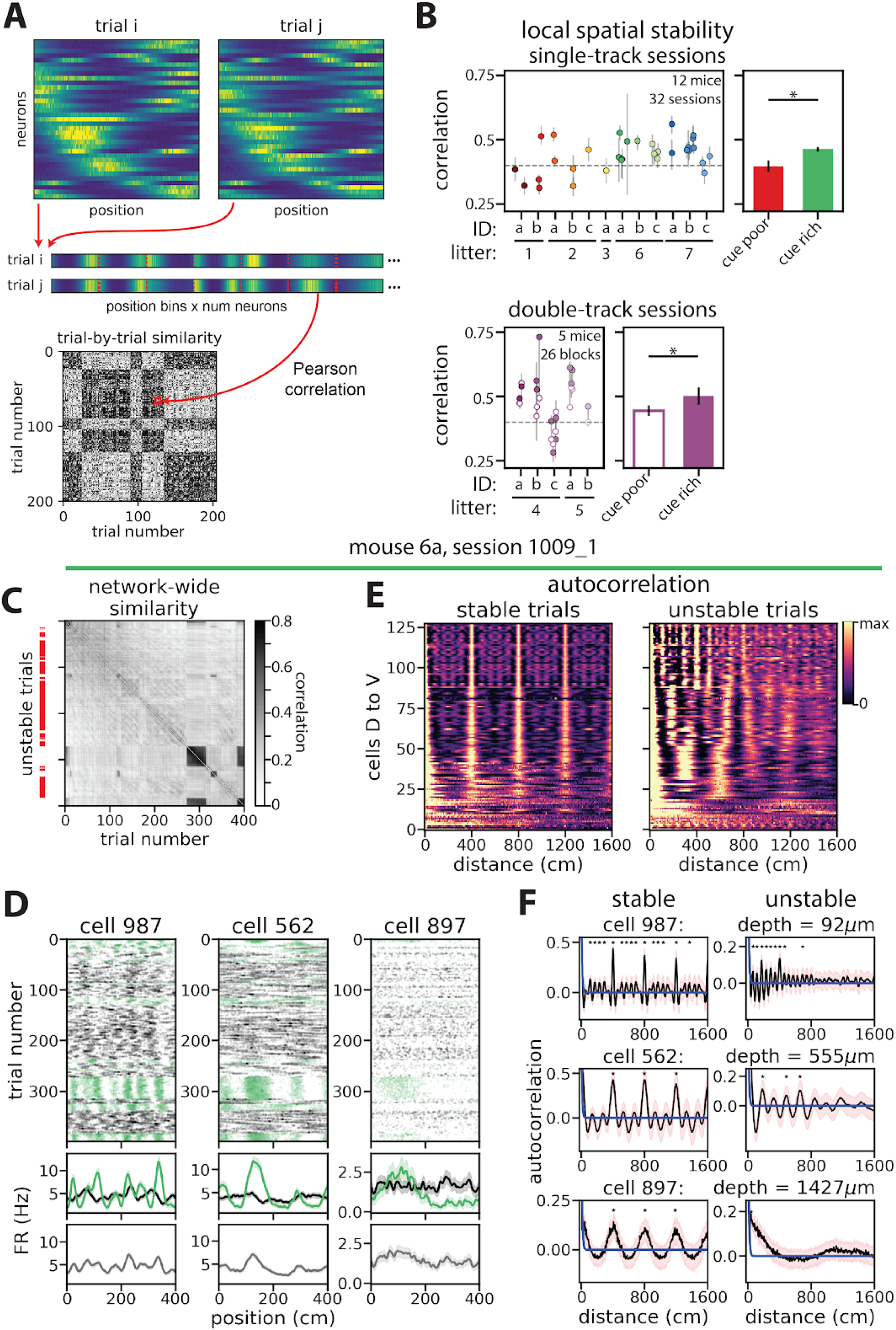
Short-term within-session stability of MEC representations. (A) For each trial, we calculated the firing rate by position for each neuron (top), linearized these matrices (middle), and computed the Pearson correlation between all trial pairs, resulting in a trial-by-trial similarity matrix of network-wide spatial coding (bottom). Each similarity matrix indicates both short-term and long-term stability in spatial coding through correlations to nearby and distant trials, respectively. (B) Short-term spatial stability, quantified as the average correlation of each trial with its four nearest neighbors, was variable across sessions recorded in a single cue rich or cue poor environment (i.e. single-track sessions, n = 32 sessions in 12 mice)(top two panels; points, single sessions; colors indicate mouse identity; gray bars, interquartile range (IQR)), as well as in alternating blocks of cue rich and cue poor tracks (i.e. double-track sessions, n = 24 blocks across 12 sessions in 5 mice)(bottom two panels; filled circles, cue rich blocks; open circles, cue poor blocks; gray bars, full session IQR)(mean short-term stability range: 0.18 to 0.73). Short-term coding stability in cue poor trials was lower than in cue rich trials for contiguous blocks of trials recorded in a given double-track session (mean short-term stability ± SEM: cue rich = 0.52 ± 0.03, cue poor = 0.45 ± 0.02; Wilcoxon two-sided signed-rank test, p = 0.028; n = 11 pairs of trial blocks in 5 mice)(rightmost panel), as well as in cue rich and poor single-track sessions recorded in different mice (mean short-term stability ± SEM: cue rich = 0.46 ± 0.01, cue poor = 0.39 ± 0.03; two-sided Wilcoxon rank-sum test, p = 0.002; n = 11 cue poor sessions in 6 mice, 21 cue rich sessions in 6 mice)(middle left). These results indicate that the number of landmarks could contribute to some of the observed variability in stability across sessions. While short-term stability was internally consistent in some sessions (n = 9 sessions and 5 blocks with IQR < 0.06), it was variable in other sessions (n = 9 sessions and 10 blocks with IQR > 0.1), suggesting that MEC coding can shift (i.e. remap) between stable and unstable regimes (within session IQR min to max: 0.039 to 0.394; n = 32 sessions in 12 mice, 24 blocks across 12 sessions in 5 mice). (C) A network-wide similarity matrix from an example session with variable short-term stability (mean local stability = 0.42, min = 0.28, max = 0.75) illustrates remapping between stable and unstable periods (red bars, trials with local stability < 0.4, i.e. “unstable trials”). (D) Single cells from the session shown in (C) have spatial firing fields in stable trials (green), but do not have position-aligned firing fields in unstable periods (black). This distinction is lost by averaging position-aligned tuning over the full session (bottom). (E, F) To examine periodic firing, we divided trials into stable and unstable periods, computed firing rate in 2 cm position bins, and computed the spatial autocorrelation for each cell’s firing. We then compared the peaks of this autocorrelation to a null model to identify cells with significant spatial periodicity (F; blue curve, null model; black curve, observed; pink shading, 95% confidence intervals)(Methods). In both stable and unstable regimes, cells exhibited spatially periodic firing that increased in period from dorsal to ventral, consistent with known grid cell properties (Brun et al., 2008; Fyhn et al., 2008; Hafting et al., 2005)(128/383 cells had spatially periodic firing). In stable regimes (short-term stability ≥ 0.4)(E, F left), spatially periodic firing aligned with the 400 cm track length across cells, as evidenced by the higher autocorrelation peaks at 400 cm, indicating that periodic spatial firing was anchored to the track landmarks in these cells. These cells maintained their spatial periodicity in unstable regimes (E, F right, ordering is the same as left panels), but spatial periodicity was not anchored to the available landmarks in these regimes; rather, each cell maintained its own preferred spatial period. Thus, transitions between spatially stable and unstable regimes may represent shifts between landmark-based and landmark-free navigational coding strategies. In panels B and F, * indicates p < 0.05.

**Fig. S3:**
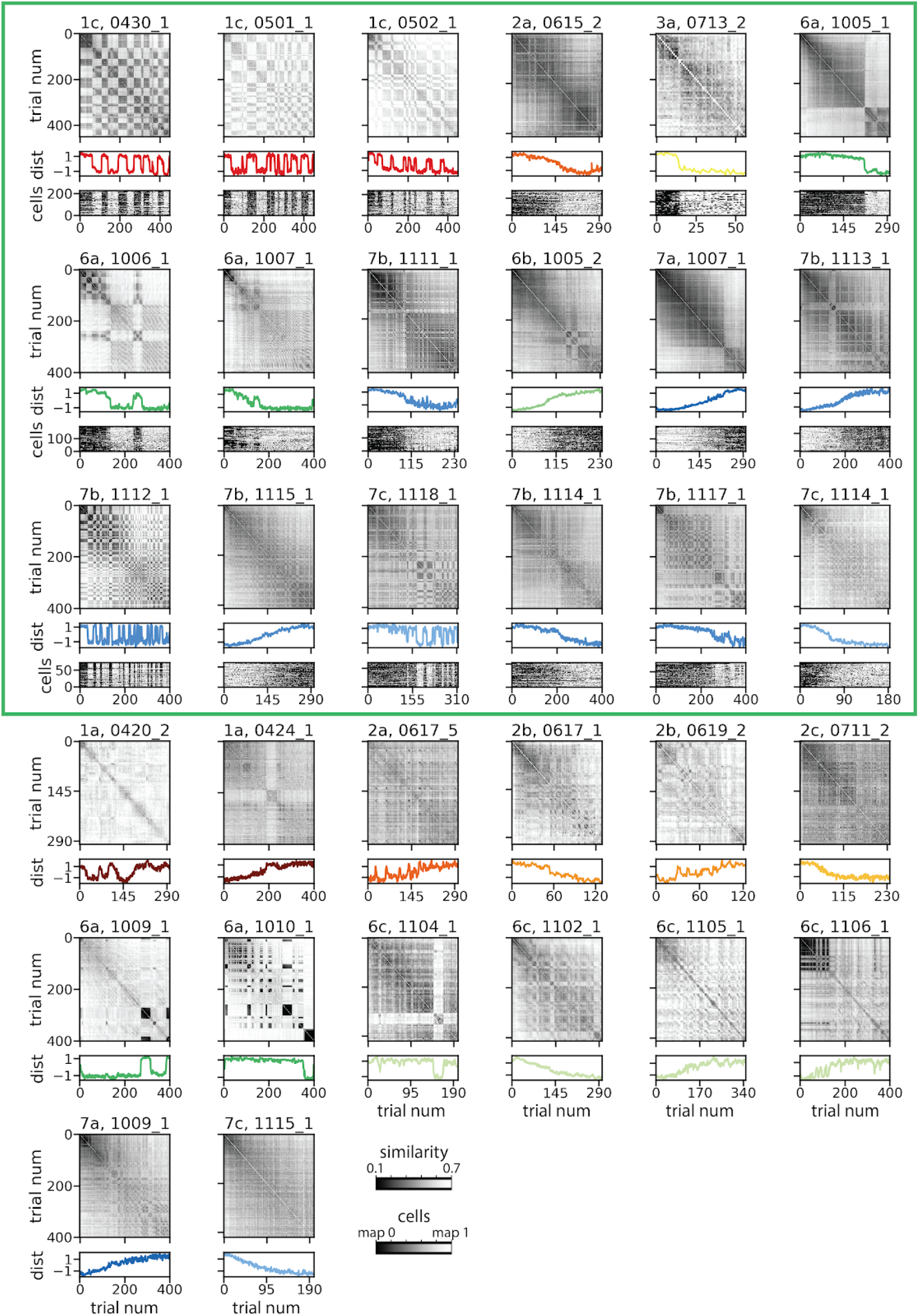
Similarity matrices and distance to 2-factor k-means cluster for all single-track sessions. Network-wide similarity matrices for many sessions showed a checkerboard pattern, indicating synchronous remapping between distinct spatial representations (top panels; colormap indicates trial-by-trial spatial correlation; black, correlation = 0.7; white, correlation = 0.1). A 2-factor k-means model (middle panels) fit sessions with variable degrees of accuracy (1 indicates in map 0 cluster centroid, −1 indicates map 1 centroid). We classified 18 sessions in 8 mice as “2-map sessions” (green box) in that they were well-described by a 2-factor k-means model (fig. 2e, green points; performance gap with PCA < 70% relative to shuffle, > 0.38). The similarity matrices from these sessions often alternated between internally stable, distinct maps (9 leftmost 2-map sessions). The distance to k-means-identified cluster qualitatively matched these transitions in most sessions. In some cases, the network appeared to transition more gradually between the two maps (9 rightmost 2-map sessions). Single cell spatial representations tended to tightly occupy each map. Heat maps of distance to cluster for each cell across all trials show discrete single-cell transitions between maps that agree precisely with the network-level remapping events (2-map sessions, bottom panels; black = in or beyond map 0 cluster centroid; white = in or beyond map 1; gray = midway between maps; cells are sorted from dorsal (top) to ventral (bottom)).

**Fig. S4:**
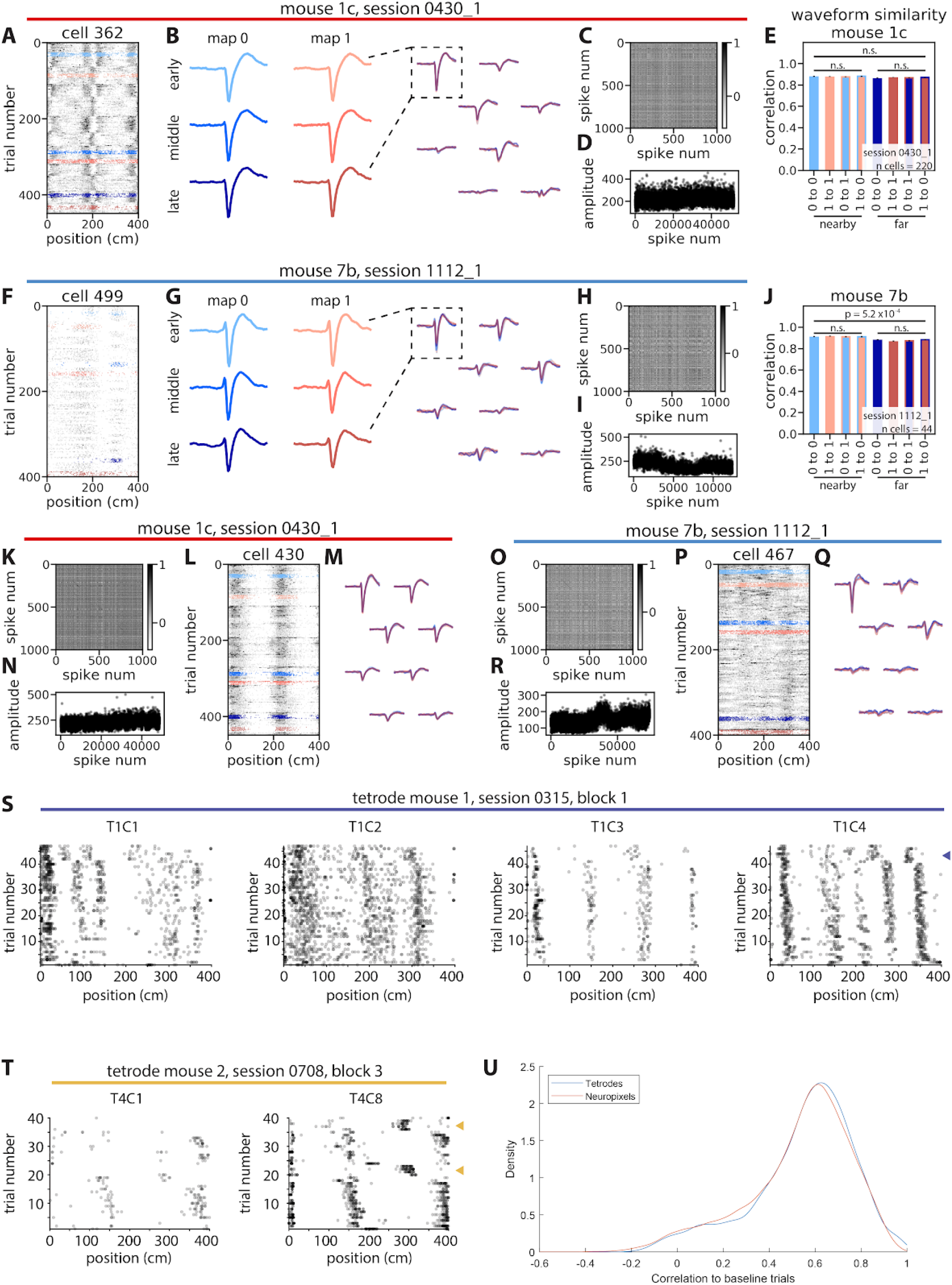
Remapping is unlikely to be an artifact of recording technique. (A-R) To account for possible artifacts of probe movement or multi-unit activity, we examined the waveforms for spikes labeled as belonging to a single unit across remapping events in two example sessions. (A-E) Illustrate that waveforms were similar across remapping events in an example cue poor session. (A) We sampled 100 spikes each from blocks of 10 trials located early (light colors), midway through (neutral colors), or late (dark colors) in the session from each map (denoted by blue and pink). (B) We then computed the average waveform across these spikes for each trial block. Example waveforms from a single contact site (right, dashed box), but calculated from different trial blocks (left; colors match A), are qualitatively similar, which holds true across channels (right; 8 best channels, overlaid; colors match A). (C) A spike-by-spike similarity matrix for this example cell compares waveforms for 1000 randomly selected spikes from throughout the session and is largely unstructured; differences in waveforms seemed not to systematically vary across remap events. (D) The amplitude of each spike was variable across the session, but these variations were not aligned with remap events. (E) Across all cells for this example session (n = 220 cells), the correlation between pairs of average waveforms was no different whether these pairs were from the same map (single color bars) or from different maps (two-color bars)(Kruskal-Wallis H-test; p = 0.11 across all pairs, n = 8 waveform average pairs from 220 cells; p = 0.96 across pairs from nearby trial blocks, n = 4 waveform average pairs; p = 0.99 across pairs from distant trial blocks, n = 4 waveform average pairs). To control for time in the session, we compared waveforms from early and midway through the session (i.e. “nearby” pairs; light colors) and waveforms from early and late in the session (i.e. “far” pairs; dark colors)(Methods). (F-J) As in (A-E), but for an example cue rich session. Waveform shapes changed slightly over the course of this session (e.g. amplitude difference in G), resulting in differences in correlation between nearby versus far pairs (Kruskal-Wallis H-test; p = 5.2×10^−4^ across all pairs, n = 8 waveform average pairs from 44 cells); however, this difference was abolished when time in session was controlled (p = 0.81 across pairs from nearby trial blocks, n = 4 waveform average pairs; p = 0.57 across pairs from distant trial blocks, n = 4 waveform average pairs)(colors match A). (K, O) As in (C, H); (L, P) as in (A, F); (M, Q) as in (B, G, right); (N, R) as in (D, I), but for an additional example cell from each session. (S-U) Tetrode data shown here was previously published (Campbell et al., 2018). In this tetrode data set, VR gain manipulations were performed (i.e. mismatch between visual and locomotor cues). To control for the possibility that frequent gain manipulations could have a lasting impact on the network’s propensity to remap, we compared the tetrode data to Neuropixels data from mice that had experienced gain manipulations (Campbell et al., 2020). All data examined for this figure are from “baseline trials” in which no gain manipulation occurred. (S, T) Cells that were co-recorded using tetrodes from two sessions. Remapping of single neurons appeared synchronized across cells and was qualitatively similar to the remapping that we observed in single Neuropixels units (e.g. Fig. 1M-Q)(arrowheads, remaps). As a measure of single cell remapping, we selected cells with stable spatial coding on the first 6 recorded trials (mean across-trial peak cross-correlation > 0.5) and compared spatial firing on the subsequent 14 trials to these baseline trials (Methods). We would expect a low cross-correlation for trials where the spatial tuning remapped from the baseline spatial map. (U) The distributions of cross-correlations for both tetrode and Neuropixels recordings were qualitatively similar, with heavy tails towards lower correlation values. Neuropixels spatial correlations were slightly lower than tetrode correlations (mean correlation to baseline ± SEM: tetrode recordings = 0.55 ± 0.003, n = 296 cells across 4144 trials, Neuropixels recordings = 0.53 ± 0.001, n = 3075 cells across 43,050 trials; two-sided Wilcoxon rank-sum test, p = 2.5×10^−6^), indicating slightly more single cell remapping in Neuropixels recordings. Note, however, that tetrode recordings were made on a track with more salient landmarks and with clearly delineated trial boundaries compared to the track used for the Neuropixels recordings, which could possibly account for this small difference.

**Fig. S5:**
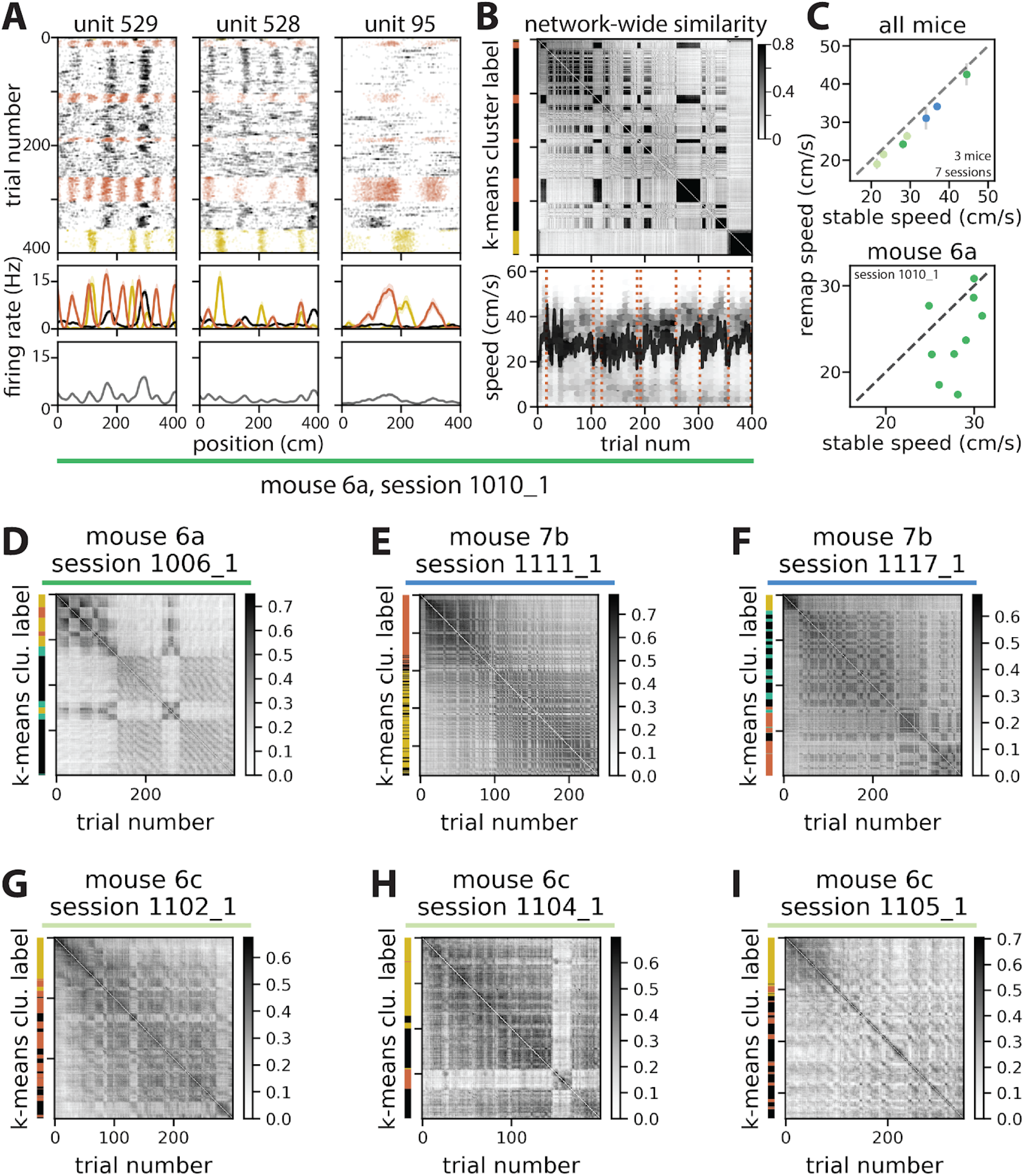
Sessions with more than 2 stable maps show many of the same characteristics as 2-map sessions. (A) (top row) Rasters for example neurons from a 3-map session, colored by k-means cluster label. Neurons exhibited distinct tuning curves in each map (middle row), while averaging neural activity over the entire session obscured this structure (bottom row)(compare to Fig. 2F). The example cells in (A) illustrate that, as in 2-map sessions, we observed cases of rate remapping and global remapping (often a mix of both) across each of the multiple maps. (B, top) The k-means assigned cluster labels (left) qualitatively matched the checkerboard structure visible in the network-wide trial-by-trial similarity matrix (right) (colorbar, spatial correlation). (B, bottom) Running speed by trial (black, trial average; gray, density) compared to remap events (dotted lines) for an example 3-map session (compare to Fig. 5A). (C, top) Similar to what we observed in 2-map sessions, for all 3- and 4-map sessions, the animal’s average running speed on remap trials was lower compared to its average running speed in the preceding stable block (mean percent difference in running speed ± SEM: 6.3 ± 2.2%; Wilcoxon two-sided signed-rank test, p = 0.0045; n = 82 remap trial/stable block pairs; “remap trials” and “stable blocks” were defined as in Fig. 4)(points, individual sessions; colors indicate mouse identity; gray bars, SEM). (C, bottom) Running speed on remap trials vs. stable blocks for an example session (points, stable block/remap trial pairs; n = 8 pairs). (D-I) As in (B, top), but for additional example 3-map (E, G, H, O) and 4-map (D, F) sessions. Note that (D-F) also met our criteria for “2-map” sessions (fig. 2–5), but additional features of the neural activity could be captured by 3-factor (E) or 4-factor (D, F) k-means models. Importantly, the relationship between running speed and remapping was preserved for these sessions regardless of model choice. N = 7 sessions in 3 mice.

**Fig. S6:**
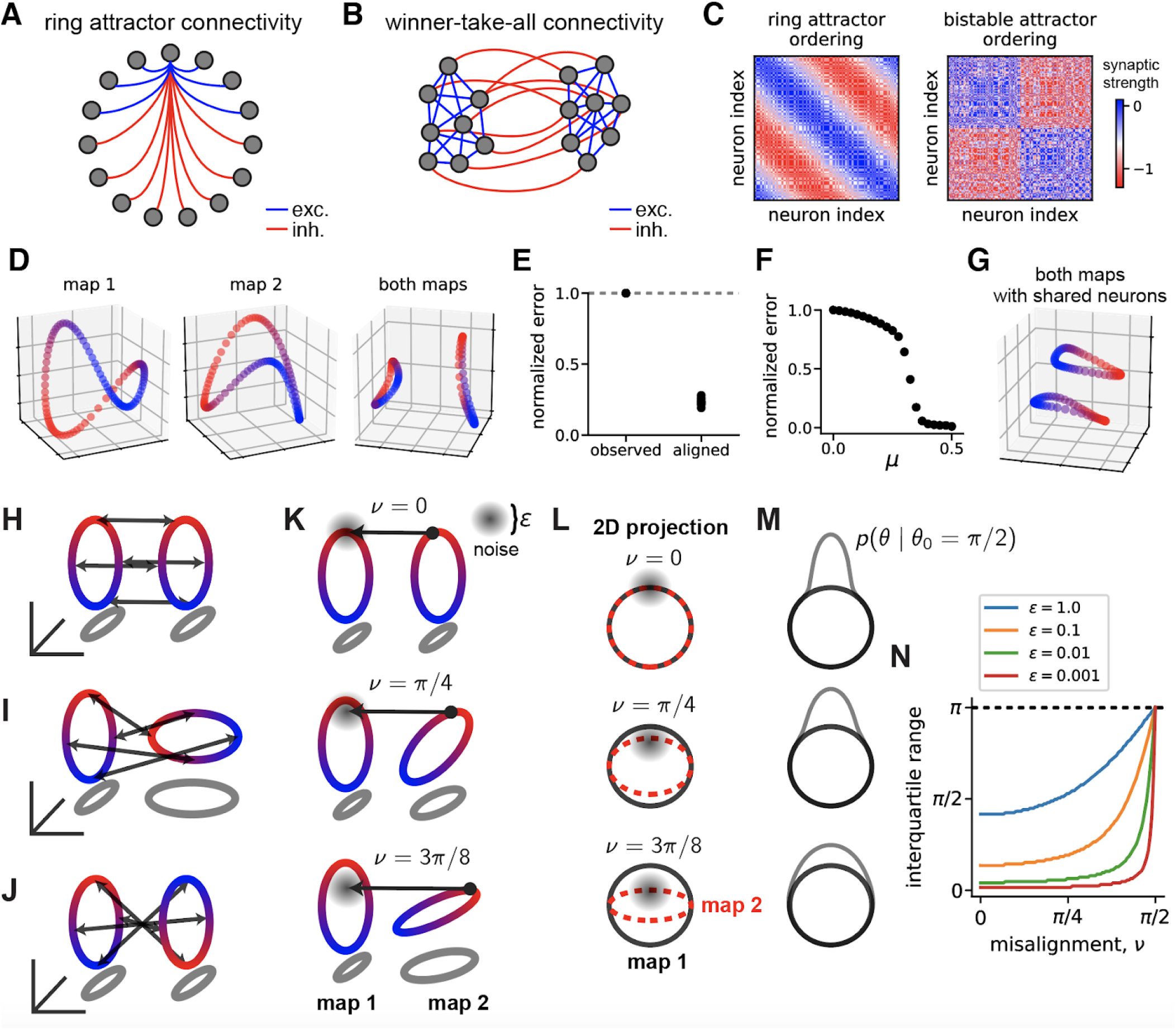
Neural circuit model supporting bistable ring attractor manifolds. (A) Illustration of ring attractor connectivity pattern (see equation S.2). Blue indicates excitatory (exc.) connections and red indicates inhibitory (inh.) connections. (B) Illustration of winner-take-all connectivity between two sub-populations of neurons (see equation S.3). Color coded as in (A). (C) Full connectivity matrix of the model. Two permutations of the neurons are shown, **W**_*ij*_ and **W**_*σ*(*i*),*σ*(*j*)_ which respectively reveal the ring attractor pattern and winner-take-all connectivity pattern. (D) PCA embeddings of the two stable ring manifolds. The blue-to-red coloring along the manifold corresponds to the encoded position, as in Figure 3 in the main text. Left, embedding of map 1 alone. Middle, embedding of map 2 alone. *Right*, simultaneous embedding of both maps. (E) Normalized root-mean-squared error of manifolds before (“observed”) and after (“aligned”) alignment by Procrustes analysis. The error is normalized to equal one for randomly rotated manifolds (dashed line); thus, we see that the simple superposition of ring and winner-take-all connectivity structures produces randomly aligned manifolds. (F) Normalized root-mean-squared error of manifolds (calculated as in panel E), as a function of the proportion of shared neurons, *μ*. (G) Example PCA embedding of a model with shared neurons (*μ* = 0.2); demonstrating better alignment than the model without shared neurons. (H) Schematic illustration of coplanar and aligned ring manifolds. Blue to red coloring of the ring manifolds corresponds to encoded position, as in Figure 3. Black double-sided arrows denote matching positions on the two manifolds, and thus correspond to appropriate remapping dimensions if the encoded position is preserved across remap events. (I) Same as panel A, but for misaligned manifolds on non-parallel planes. Note that remapping dimensions are no longer parallel to each other and depend on the location along the manifold. (J) Same as panel A, but with the manifolds misaligned by a vertical reflection. As in panel B, remapping dimensions are no longer parallel. (K) Schematic illustration of remapping (black arrow) in the presence of noise (gray spherical blur) and three different levels of rotational misalignment. Map 2 is rotated *v* radians away from a plane parallel to map 1. We consider remapping from an initial position *θ*_0_ = π/2 0n map 2. (L) Projection of 3D plots in panel D onto the plane spanned by map 1. Map 1 is shown in black; the projection of map 2 is shown as a dashed red line. For nonzero *v* the projection of map 2 onto this 2D plane is an ellipse. The remapping noise is a bivariate Gaussian distribution after this projection (gray circular blur). (M) Assuming attractor dynamics project the distribution of activity to the nearest point on map 1, the final position along the ring manifold, *θ*, follows a projected normal distribution. The density function of this distribution is plotted in gray for the three levels of misalignment shown in panel E. (N) The interquartile range of the projected normal distribution is plotted as a function of *v* for four levels of noise, *ϵ*. Uncertainty in the final position, *θ*, increases as the noise increases and as the misalignment increases.

## Supplementary Discussion

### Attractor Models of Bistable Spatial Maps

Attractor networks are a class of models that describe how neural circuits can create persistent internal representations about the state of the world, even after sensory inputs are removed (Amari, 1977; Samsonovich and McNaughton, 1997; Seung, 1996). Attractor models of MEC dynamics were implemented soon after the discovery of grid cells (Burak and Fiete, 2009; Fuhs and Touretzky, 2006; Guanella et al., 2007; McNaughton et al., 2006) and have influenced subsequent experimental research (Couey et al., 2013; Pastoll et al., 2013; Stensola et al., 2012; Yoon et al., 2013). In their simplest form, these models construct a single attractor manifold, corresponding to a single internal spatial map. Our observation that MEC spontaneously remaps between multiple internal representations reveals one way in which neural activity is more complex than this idealized model. In this section, we show how to reconcile our major experimental findings with existing theory.

Attractor models are often formulated as a system of differential equations describing neural firing rates:

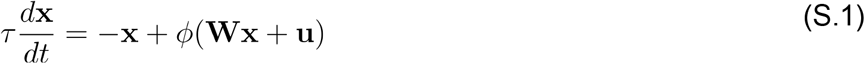

where **x** ∈ ℝ^*N*^ denotes a vector of firing rates for a network containing *N* neurons, **W** ∈ ℝ^*N* × *N*^ is the “connectivity matrix” holding synaptic weights, *ϕ*(·) is an elementwise nonlinear activation function, **u** ∈ ℝ^*N*^ is a constant input to the network, and *τ* > 0 is the time constant of the system. We assume that *ϕ*(*x*) = 1/(1 + exp(−x)), though other choices of activation function are possible. More detailed models have been developed with greater attention to biological plausibility (Laing and Chow, 2001; Navratilova et al., 2012; Widloski and Fiete, 2014); however, these share many of the same fundamentals as the simpler rate model in eq. (S.1).

Values of **x** which satisfy *d***x**/*dt* = 0 are called *fixed points*. Each fixed point represents a persistent firing rate configuration of the network. Of particular interest are *attractive* fixed points (also called asymptotically stable fixed points): a fixed point **x**_∗_ is attractive if the dynamical trajectory **x**(*t*) approaches in the **x**_∗_ limit as *t* ⟶ ∞ after being initialized at **x**(0) = **x**_∗_ + *δ* where *δ* denotes a suitably small perturbation. *Attractor manifolds* can be thought of as continuous sets of attractive fixed points. While attractive fixed points are limited to representing a discrete set of states, attractor manifolds can represent continuous quantities, such as the orientation of a visual stimulus (Ben-Yishai et al., 1995), motor neuron drive (Seung, 1996), the animal’s heading direction (Skaggs et al., 1995), or position in space (Burak and Fiete, 2009; Samsonovich and McNaughton, 1997).

In our experiments, mice traversed a 1-dimensional virtual hallway that seamlessly looped back to the starting position after 400 cm. This can be thought of as a virtual reality analogue to a circular maze environment. The attractor manifold framework predicts that neural activity mirrors the structure of this external environment—specifically, the fixed points of the network are arranged in a 1-dimensional ring attractor manifold. Further, the velocity of the animal on the track is calibrated to the velocity in neural firing rate space along the ring attractor so that every physical location on the track is one-to-one matched to a position along the ring. While previous studies have used ring attractor networks to model subpopulations of comodular grid cells within MEC (Burak and Fiete, 2009; Giocomo et al., 2011), all of our analyses and models consider the full MEC network including landmark cells, border cells, object vector cells, and cells with mixed or unknown selectivity. Our approach is not necessarily incompatible with models that restrict their focus to sub-structures within the circuit.

Ring attractor networks have been proposed in a variety of contexts (Ben-Yishai et al., 1995; Hansel and Sompolinsky, 1998; Skaggs et al., 1995; Zhang, 1996). Most of these models utilize a connectivity pattern in which neurons are arranged in a ring with short-range excitatory connections and long-range inhibitory connections (fig. S6A). Intuitively, this creates a stable “bump” of activity at one location in the ring. Neurons encoding the animal’s velocity can then move the position of this bump (Navratilova et al., 2012); however, for our purposes, it is unnecessary to model these details explicitly.

The notion that a single network may store multiple attractors, corresponding to different internal maps, has been previously considered in several papers (Romani and Tsodyks, 2010; Roudi and Treves, 2008; Samsonovich and McNaughton, 1997; Stringer et al., 2004). Following these works, we set the network’s connectivity to be a linear superposition of attractor networks---that is, to store attractor manifolds we set 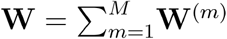, where each **W**^(*m*)^ ∈ **W**^*N* × *N*^ holds the connectivity pattern of a single map. If *M* ≪ *N*, and if the maps are sufficiently decorrelated (e.g. if the neuron indices are randomly permuted for each **W**^(*m*)^), then the attractors can operate independently (Samsonovich and McNaughton, 1997).

There are potentially multiple models within this framework that could be used to account for the remapping events we experimentally observed. Here, we outline a simple possibility that combines a classic ring attractor connectivity pattern (fig. S6A), with a “winner-take-all” connectivity pattern (fig. S6B). Intuitively, the winner-take-all pattern creates two sub-populations of neurons that mutually exclude each other from firing. Because the ring attractor structure is present within both sub-populations, two mutually exclusive spatial maps are created. To instantiate the model numerically, we define a fine grid over ring angles, *θ*_*i*_ = 2*πi*/*N* for *i* ∈ {1, …, *N*}. Additionally, for each neuron we define a sub-population indicator variable 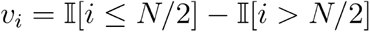; where 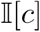 is a binary indicator function which evaluates to one if *c* is true and zero if *c* is false. Then, let *σ*(·) denote a random permutation of the *N* neurons, and define the connectivity of the network to be **W** = *W*^(1)^ + *W*^(2)^, where

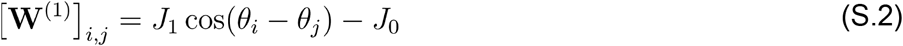

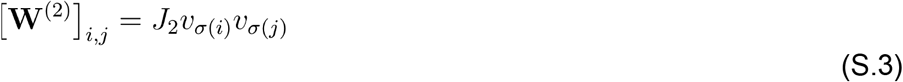

The connectivity encoded in *W*^(1)^ implements the ring attractor, while *W*^(2)^ implements the winner take all connectivity. The three scalar hyperparameters, *J*_0_ > 0, *J*_1_ > 0, *J*_2_ > 0, respectively determine the strength of a global inhibition term, the strength of connections modulated by the ring, and the strength of the winner-take-all connectivity. For now, we assume the network receives no input, **u** = 0.

Figure S6C shows *W*^*i,j*^ and *W*^*σ(i)*,*σ(j)*^ for a small network with *N* = 150 neurons; this visualization demonstrates that simply re-ordering the neurons is sufficient to reveal the two connectivity patterns embedded in the same network. We numerically simulated a network with *N* = 1000 neurons to explore whether this model produced the expected attractor structures. For these simulations we set *J*_0_ = *J*_1_ = 0.01 and *J*_2_ = 0.004. The steady-state of eq. (S.1) is solely determined by the initial state, **x**(0), which we set equal to

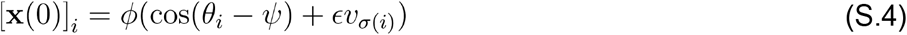

where *ψ* ∈ [0, 2π) is an angle specifying the tuning of initial activity along the ring, and *ϵ* scales the competitive winner-take-all connectivity pattern. When *ϵ* > 0 the network activity reliably converges to one of the spatial maps; conversely, when *ϵ* < 0, the network converges to the other map. As *ψ* is varied and *ϵ* is kept fixed, the steady-state activity traces out the expected ring attractor manifold (fig. S6D).

In contrast to the experimental data, when both maps are simultaneously embedded into the same 3D space by PCA, the ring manifolds do not appear aligned (fig. S6D, right). We confirmed this by using Procrustes analysis to quantify manifold alignment (as done for the experimental data in fig. 4). Indeed, measured relative to a shuffle control, the root-mean-squared-error (RMSE) substantially decreased after the optimal rotational alignment was applied to the manifolds (fig. S6E). Overall, this suggests that the natural alignment of the ring manifolds is a non-trivial feature of the experimental data that is not universally present in attractor models.

While the attractor model defined by equations (S.2 - S.3) does not capture the experimentally observed alignment of the spatial manifolds, it is relatively straightforward to incorporate remapping into the model. To do this, we introduce a noise term (formally, Brownian motion) into the dynamics of equation (S.1) and numerically integrate the dynamics by the Euler–Maruyama method. Inspired by the experimental findings in Fig. 4K-L, we use a mix of isotropically distributed noise with occasional perturbations preferentially oriented in the dimension separating the manifolds. These directed perturbations occasionally push the network activity from one ring attractor to the other, resulting in a remap event (Supplemental Video 1).

Finally, we investigated whether the model could be modified to produce aligned ring manifolds, as we experimentally observed. Indeed, introducing a population of “shared neurons” that participated in both spatial maps was sufficient to reproduce this feature of the data. We implemented this model extension by defining

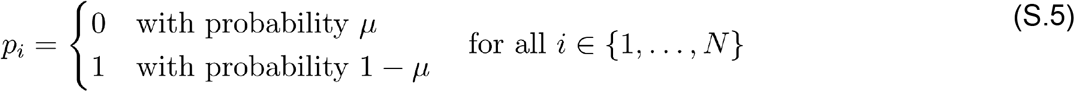

where *μ* ∈ [0, 1] corresponds to the proportion of “shared neurons.” Then, we modified equation (S.3) to be:

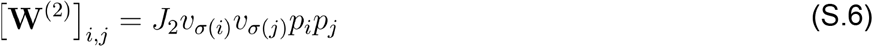

and set the network input to **u**_*i*_ = *J*_4_(1 − *p*_*i*_) for each neuron. Intuitively, *J*_4_ scales additional excitatory input to the “shared neurons” that lack the mutual excitation from the winner-take-all connectivity. We set *J*_4_ = 4 in our numerical simulations. As *μ* was increased, we found the two maps were pulled closer together and became more aligned, eventually fusing into a single ring manifold (fig. S6F). At intermediate values, the rings appear geometrically aligned in PCA embeddings (fig. S6G), but are still capable of operating independently. Supplemental Video 2 demonstrates this, showing that the model exhibits remapping events akin to those described above for the initial simulations without shared neurons (the parameters of the noise process needed no modification for this simulation).

Overall, these results outline a model of our experimental results in which directed noise spontaneously induces remapping in a network combining winner-take-all and ring attractor connectivity motifs. This provides additional mathematical rigor and precision to substantiate the conceptual model in figure 4N, and also lays the foundation for future modeling work. A key open question is how running speed—and, more broadly, behavioral state—should be modeled. These factors tend to increase the probability of remapping in our experimental data (see fig. 5). In the model similar effects might be accomplished in several ways, such as modulating the scale of the noise term, manipulating the input to the network, or by incorporating neuromodulatory effects into the model. Alternative mechanisms for maintaining multiple maps, such as correlated ring attractor manifolds (Romani and Tsodyks, 2010), might also be investigated. Finally, the attractor model presented here does not account for the reality that many MEC neurons have multiple spatial firing fields, and does not consider the possibility of more sophisticated interactions between landmark and grid cells (Campbell et al., 2018; Kang and Balasubramanian, 2019; Ocko et al., 2018). Nonetheless, it is noteworthy that the simple and preliminary model outlined here is sufficient to recapitulate most of our experimental observations.

### Consequences of Manifold Alignment For Decoding and Remapping

As mentioned in the main text, the geometrical alignment of the spatial manifolds has two advantageous consequences. First, it implies that simple linear decoders of MEC’s representation of position can be robust to remapping dynamics. Second, it ensures that a simple remapping mechanism—namely, bistable attractor dynamics combined with variability in neural firing—preserves the network’s representation of position and is noise tolerant. In this section, we briefly formalize these two conclusions.

Recall our notation where 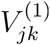 and 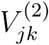 respectively denote the first and second spatial manifolds, with *j* ∈ {1, ∈, *J*} indexing position bins, and *k* ∈ {1, ∈, *K*} indexing neurons. In the main text, we empirically estimated 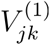 and 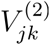 by applying k-means clustering to experimental data. Here, we treat the problem on more general terms and assume that the manifolds are known.

The first result—that manifold alignment enables robust linear decoding—is easy to demonstrate. Intuitively, for aligned ring manifolds, remapping is accomplished by translation along a single dimension (fig. S6H), while more complex and position-dependent remapping dimensions are required for misaligned rings (fig. S6I-J). Suppose that the two manifolds are perfectly aligned, i.e., there exists some translation vector *z*_*k*_ for which:

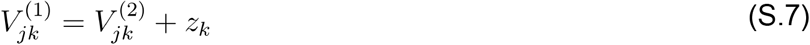

Then, let *w*_*k*_ denote a set of regression weights associated with some linear decoder of position (see *Methods*; though two linear features are required since the dependent variable is a circular quantity, the reasoning applies equally well to each feature vector). This linear feature is insensitive to remapping if Σ_*k*_*w*_*k*_*z*_*k*_ = 0, since this implies:

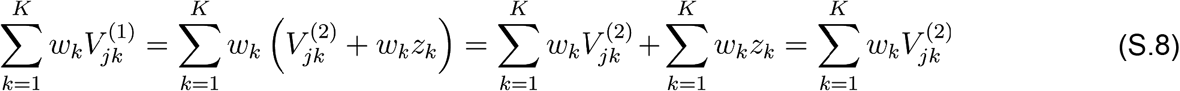

Thus, so long as the remapping dimension is orthogonal to the regression weights, any decoder making use of this linear feature will perform equally well in each map. This is only a minor constraint as it leaves *N* − 1 orthogonal dimensions for the linear decoder to utilize.

Constructing a robust decoder with fixed weights is more difficult when the two manifolds are misaligned. Suppose, for simplicity, that 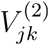 is centered at the origin (i.e. 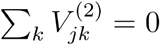 for each position bin indexed by *j*). Further suppose that the two manifolds are related by an orthogonal matrix *Q*_*kk′*_ in addition to a translation:

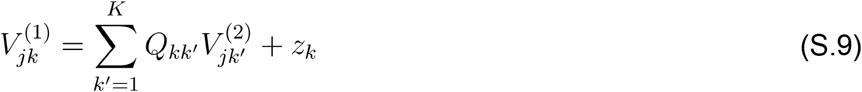

Then, even if the regression weights were orthogonal to the remapping dimension, we would additionally require that Σ_*k′*_*Q*_*kk′*_w_*k′*_ = *w*_*k*_ in order to obtain an identical linear feature. Geometrically, this corresponds to *w*_*k*_ being a fixed axis of rotation or reflection (equivalently, *w*_*k*_ is an eigenvector *Q*_*kk′*_ with an eigenvalue equal to one). Some orthogonal transformations have no such fixed axes, so in general we cannot expect to find a linear decoder that works equally well in each map if the manifolds are misaligned.

In addition to enabling simple decoding strategies, the alignment of the ring manifolds also simplifies intrinsic circuit mechanisms for remapping. This second result shares many intuitions with the above discussion of decoders. Namely, if the manifolds are aligned (i.e. the relation S.7 holds) then only a single remapping dimension is required (fig. S6H). Thus a perturbation to network activity along a fixed dimension (corresponding to *z*_*k*_ in equation S.7) is a viable remapping mechanism. Note that this model of remapping is entirely agnostic to the animal’s position, which is not generally sensible for the case of misaligned manifolds (fig. S6I-J).

Incorporating noise into each remapping event further complicates the case of misaligned manifolds. Indeed, if we make several simplifying assumptions (which could be relaxed by future work), it is possible to analytically characterize the error in positional coding induced by remapping as a function of manifold misalignment. We assume that the two ring manifolds are circular and embedded in 2-dimensional planes, and we model remapping as a projection of the activity from one map (arbitrarily labeled map 2) onto the other map (labeled map 1). We add an isotropic Gaussian noise term with covariance *ϵ***I** to model instability in the remapping mechanism. Figure S6K schematically illustrates this model as map 2 is rotated to be non-parallel with map 1. Let *v* denote the angle of rotation, such that *v* = 0 corresponds to the case where the manifolds are aligned. It is evident that the projection of map 2 onto map 1 is equivalent whether the rotation is clockwise or counterclockwise rotations, so we can restrict ourselves to considering *=* ∈ [0, *π*]. Further, we are principally interested in the interval *=* ∈ [0, *π*/2], since for *=* > *π*/2 the manifolds are irreparably misaligned (see fig. S6J) and would require a more complex, position-dependent remapping mechanism. The distortions and uncertainty introduced by this noisy remapping process depend on the initial position of neural activity on map 2. For simplicity, we focus on the case where this initial position is located at the “top” of map 2 (as illustrated in fig. S6K), and leave a more comprehensive analysis to future work.

After projecting the location of neural activity onto the plane containing map 1 and adding isotropic Gaussian noise, we assume that the ring attractor dynamics move the activity to the closest point on the manifold defining map 1. This final step can be understood as projecting a bivariate Gaussian density onto the unit circle. This results in the *projected normal distribution*, which has been characterized in the circular statistics literature and has a closed form density function (Wang and Gelfand, 2013). When the covariance of the projected density is diagonal (as we assume), the resulting distribution is symmetric and unimodal and has a similar appearance to the more familiar von Mises distribution. Figure S6L illustrates the distribution of neural activity before projecting the bivariate density onto the circle. Figure S6M illustrates the final probability density function over the circle—notice that the width of this distribution increases as the manifolds become more misaligned, even though the scale of the noise, *ϵ*, is held fixed. The interquartile range is plotted as a function of *v* in figure S6N for various choices of *ϵ*. In the limit as *v* ⟶ *π*/2 the distribution becomes uniform on the circle, representing a complete loss of position coding from map 2. Interestingly, noise has a much smaller effect for smaller misalignments ((*v* < *π*/4)), relative to manifolds with large misalignments (*v* > *π*/4), suggesting that the simple position-independent remapping scheme may be naturally tolerant to some imperfections.

## Supplemental Video Legends

**Video S1:** Model of bistable ring attractor dynamics in the presence of noise. A 3-dimensional PCA embedding of the two ring attractors, with red-to-blue circular color scheme denoting position, is shown as in fig. S6D (right panel). The moving black dot represents the evolution of circuit activity through neural firing rate space in the presence of noise (Wiener process with identity covariance); the gray line shows the recent trajectory. Time units are arbitrary. Occasional noise perturbations restricted the dimension separating the two manifolds cause remapping in a probabilistic manner.

**Video S2:** Model of bistable ring attractor dynamics with shared neurons. The setup is the same as Video S1, but with 20% shared neurons between each map (*μ* = 0.2). This minor modification is sufficient to provide the visual appearance of aligned rings in closer agreement to the biological data analyzed in the main text.

## References

Allen Institute for Brain Science (2004). Mouse Brain Atlas.

Amari, S. (1977). Dynamics of pattern formation in lateral-inhibition type neural fields. Biol. Cybern. 27, 77–87.

Bant, J.S., Hardcastle, K., Ocko, S.A., and Giocomo, L.M. (2020). Topography in the Bursting Dynamics of Entorhinal Neurons. Cell Rep. 30, 2349–2359.e7.

Barry, C., Hayman, R., Burgess, N., and Jeffery, K.J. (2007). Experience-dependent rescaling of entorhinal grids. Nature Neuroscience 10, 682–684.

Ben-Yishai, R., Bar-Or, R.L., and Sompolinsky, H. (1995). Theory of orientation tuning in visual cortex. Proc. Natl. Acad. Sci. U. S. A. 92, 3844–3848.

Boccara, C.N., Nardin, M., Stella, F., O’Neill, J., and Csicsvari, J. (2019). The entorhinal cognitive map is attracted to goals. Science 363, 1443–1447.

Brun, V.H., Solstad, T., Kjelstrup, K.B., Fyhn, M., Witter, M.P., Moser, E.I., and Moser, M.-B. (2008). Progressive increase in grid scale from dorsal to ventral medial entorhinal cortex. Hippocampus 18, 1200–1212.

Burak, Y., and Fiete, I.R. (2009). Accurate path integration in continuous attractor network models of grid cells. PLoS Comput. Biol. 5, e1000291.

Butler, W.N., Hardcastle, K., and Giocomo, L.M. (2019). Remembered reward locations restructure entorhinal spatial maps. Science 363, 1447–1452.

Calhoun, A.J., Pillow, J.W., and Murthy, M. (2019). Unsupervised identification of the internal states that shape natural behavior. Nat. Neurosci. 22, 2040–2049.

Campbell, M., Attinger, A., Ocko, S., Ganguli, S., and Giocomo, L. (2020). Bayesian inference through attractor dynamics in medial entorhinal cortex. In Poster Session 3, pp. III – 95.

Campbell, M.G., Ocko, S.A., Mallory, C.S., Low, I.I.C., Ganguli, S., and Giocomo, L.M. (2018). Principles governing the integration of landmark and self-motion cues in entorhinal cortical codes for navigation. Nat. Neurosci. 21, 1096–1106.

Chatfield, C. (1984). The Analysis of Time Series: An Introduction.

Churchland, M.M., Cunningham, J.P., Kaufman, M.T., Foster, J.D., Nuyujukian, P., Ryu, S.I., and Shenoy, K.V. (2012). Neural population dynamics during reaching. Nature 487, 51–56.

Couey, J.J., Witoelar, A., Zhang, S.-J., Zheng, K., Ye, J., Dunn, B., Czajkowski, R., Moser, M.-B., Moser, E.I., Roudi, Y., et al. (2013). Recurrent inhibitory circuitry as a mechanism for grid formation. Nat. Neurosci. 16, 318–324.

Diehl, G.W., Hon, O.J., Leutgeb, S., and Leutgeb, J.K. (2017). Grid and Nongrid Cells in Medial Entorhinal Cortex Represent Spatial Location and Environmental Features with Complementary Coding Schemes. Neuron 94, 83–92.e6.

Eckart, C., and Young, G. (1936). The approximation of one matrix by another of lower rank. Psychometrika 1, 211–218.

Elsayed, G.F., and Cunningham, J.P. (2017). Structure in neural population recordings: an expected byproduct of simpler phenomena? Nat. Neurosci. 20, 1310–1318.

Fisher, N.I., and Lee, A.J. (1992). Regression Models for an Angular Response. Biometrics 48, 665–677.

Fuhs, M.C., and Touretzky, D.S. (2006). A spin glass model of path integration in rat medial entorhinal cortex. J. Neurosci. 26, 4266–4276.

Fyhn, M., Hafting, T., Treves, A., Moser, M.-B., and Moser, E.I. (2007). Hippocampal remapping and grid realignment in entorhinal cortex. Nature 446, 190–194.

Fyhn, M., Hafting, T., Witter, M.P., Moser, E.I., and Moser, M.-B. (2008). Grid cells in mice. Hippocampus 18, 1230–1238.

Gil, M., Ancau, M., Schlesiger, M.I., Neitz, A., Allen, K., De Marco, R.J., and Monyer, H. (2018). Impaired path integration in mice with disrupted grid cell firing. Nature Neuroscience 21, 81–91.

Giocomo, L.M., Moser, M.-B., and Moser, E.I. (2011). Computational models of grid cells. Neuron 71, 589–603.

Gower, J.C., Statistics Department John C Gower, Dijksterhuis, G.B., and Consumer and Market Insight Agrotechnology and Food Innovations B V Wageningen University and Research Centre and Department of Marketing and Marketing Research Faculty of Economics Garmt B Dijksterhuis (2004). Procrustes Problems (OUP Oxford).

Guanella, A., Kiper, D., and Verschure, P. (2007). A model of grid cells based on a twisted torus topology. Int. J. Neural Syst. 17, 231–240.

Hafting, T., Fyhn, M., Molden, S., Moser, M.-B., and Moser, E.I. (2005). Microstructure of a spatial map in the entorhinal cortex. Nature 436, 801–806.

Hansel, D., and Sompolinsky, H. (1998). Modeling Feature Selectivity in Local Cortical Circuits. In Methods in Neuronal Modeling: From Synapses to Networks, C. Koch, and I. Segev, eds. (Cambridge, MA, USA: MIT Press), pp. 499–567.

Hardcastle, K., Maheswaranathan, N., Ganguli, S., and Giocomo, L.M. (2017). A Multiplexed, Heterogeneous, and Adaptive Code for Navigation in Medial Entorhinal Cortex. Neuron 94, 375–387.e7.

Harris, C.R., Jarrod Millman, K., van der Walt, S.J., Gommers, R., Virtanen, P., Cournapeau, D., Wieser, E., Taylor, J., Berg, S., Smith, N.J., et al. (2020). Array Programming with NumPy.

Hinman, J.R., Brandon, M.P., Climer, J.R., Chapman, G.W., and Hasselmo, M.E. (2016). Multiple Running Speed Signals in Medial Entorhinal Cortex. Neuron 91, 666–679.

Høydal, Ø.A., Skytøen, E.R., Andersson, S.O., Moser, M.-B., and Moser, E.I. (2019). Object-vector coding in the medial entorhinal cortex. Nature 568, 400–404.

Hulse, B.K., Lubenov, E.V., and Siapas, A.G. (2017). Brain State Dependence of Hippocampal Subthreshold Activity in Awake Mice. Cell Rep. 18, 136–147.

Hunter, J.D. (2007). Matplotlib: A 2D Graphics Environment. Comput. Sci. Eng. 9, 90–95.

Jennings, J.H., Kim, C.K., Marshel, J.H., Raffiee, M., Ye, L., Quirin, S., Pak, S., Ramakrishnan, C., and Deisseroth, K. (2019). Interacting neural ensembles in orbitofrontal cortex for social and feeding behaviour. Nature 565, 645–649.

Jones, E., Oliphant, T., Peterson, P., and Others (2001). SciPy: Open source scientific tools for Python.

Jun, J.J., Steinmetz, N.A., Siegle, J.H., Denman, D.J., Bauza, M., Barbarits, B., Lee, A.K., Anastassiou, C.A., Andrei, A., Aydın, Ç., et al. (2017). Fully integrated silicon probes for high-density recording of neural activity. Nature 551, 232–236.

Kang, L., and Balasubramanian, V. (2019). A geometric attractor mechanism for self-organization of entorhinal grid modules. eLife 8.

Kaufman, M.T., Churchland, M.M., Ryu, S.I., and Shenoy, K.V. (2014). Cortical activity in the null space: permitting preparation without movement. Nat. Neurosci. 17, 440–448.

Keene, C.S., Bladon, J., McKenzie, S., Liu, C.D., O’Keefe, J., and Eichenbaum, H. (2016). Complementary Functional Organization of Neuronal Activity Patterns in the Perirhinal, Lateral Entorhinal, and Medial Entorhinal Cortices. J. Neurosci. 36, 3660–3675.

Kentros, C.G., Agnihotri, N.T., Streater, S., Hawkins, R.D., and Kandel, E.R. (2004). Increased attention to spatial context increases both place field stability and spatial memory. Neuron 42, 283–295.

Kerr, K.M., Agster, K.L., Furtak, S.C., and Burwell, R.D. (2007). Functional neuroanatomy of the parahippocampal region: the lateral and medial entorhinal areas. Hippocampus 17, 697–708.

Knierim, J.J., Kudrimoti, H.S., and McNaughton, B.L. (1998). Interactions Between Idiothetic Cues and External Landmarks in the Control of Place Cells and Head Direction Cells. Journal of Neurophysiology 80, 425–446.

Kolda, T.G., and Bader, B.W. (2009). Tensor Decompositions and Applications. SIAM Rev. 51, 455–500.

Kriegeskorte, N., and Douglas, P.K. (2019). Interpreting encoding and decoding models. Curr. Opin. Neurobiol. 55, 167–179.

Krupic, J., Bauza, M., Burton, S., Barry, C., and O’Keefe, J. (2015). Grid cell symmetry is shaped by environmental geometry. Nature 518, 232–235.

Laing, C.R., and Chow, C.C. (2001). Stationary bumps in networks of spiking neurons. Neural Comput. 13, 1473–1494.

Marozzi, E., Ginzberg, L.L., Alenda, A., and Jeffery, K.J. (2015). Purely Translational Realignment in Grid Cell Firing Patterns Following Nonmetric Context Change. Cereb. Cortex 25, 4619–4627.

McNaughton, B.L., Battaglia, F.P., Jensen, O., Moser, E.I., and Moser, M.-B. (2006). Path integration and the neural basis of the “cognitive map.” Nat. Rev. 7, 663–678.

Muller, R.U., and Kubie, J.L. (1987). The effects of changes in the environment on the spatial firing of hippocampal complex-spike cells. J. Neurosci. 7, 1951–1968.

Munn, R.G.K., Mallory, C.S., Hardcastle, K., Chetkovich, D.M., and Giocomo, L.M. (2020). Entorhinal velocity signals reflect environmental geometry. Nature Neuroscience.

Navratilova, Z., Giocomo, L.M., Fellous, J.-M., Hasselmo, M.E., and McNaughton, B.L. (2012). Phase precession and variable spatial scaling in a periodic attractor map model of medial entorhinal grid cells with realistic after-spike dynamics. Hippocampus 22, 772–789.

Niell, C.M., and Stryker, M.P. (2010). Modulation of visual responses by behavioral state in mouse visual cortex. Neuron 65, 472–479.

Ocko, S.A., Hardcastle, K., Giocomo, L.M., and Ganguli, S. (2018). Emergent elasticity in the neural code for space. Proc. Natl. Acad. Sci. U. S. A. 115, E11798–E11806.

O’Keefe, J., and Conway, D.H. (1978). Hippocampal place units in the freely moving rat: why they fire where they fire. Exp. Brain Res. 31, 573–590.

Pachitariu, M., Steinmetz, N.A., Kadir, S.N., Carandini, M., and Harris, K.D. (2016). Fast and accurate spike sorting of high-channel count probes with KiloSort. In Advances in Neural Information Processing Systems, pp. 4448–4456.

Pastoll, H., Solanka, L., van Rossum, M.C.W., and Nolan, M.F. (2013). Feedback inhibition enables θ-nested γ oscillations and grid firing fields. Neuron 77, 141–154.

Pedregosa, F., Varoquaux, G., Gramfort, A., Michel, V., Thirion, B., Grisel, O., Blondel, M., Prettenhofer, P., Weiss, R., Dubourg, V., et al. (2011). Scikit-learn: Machine learning in Python. The Journal of Machine Learning Research 12, 2825–2830.

Pewsey, A., and García-Portugués, E. (2020). Recent advances in directional statistics.

Presnell, B., Morrison, S.P., and Littell, R.C. (1998). Projected Multivariate Linear Models for Directional Data. Journal of the American Statistical Association 93, 1068–1077.

Romani, S., and Tsodyks, M. (2010). Continuous attractors with morphed/correlated maps. PLoS Comput. Biol. 6.

Roudi, Y., and Treves, A. (2008). Representing where along with what information in a model of a cortical patch. PLoS Comput. Biol. 4, e1000012.

Rule, M.E., Loback, A.R., Raman, D.V., Driscoll, L., Harvey, C.D., and O’Leary, T. (2020). Stable task information from an unstable neural population. bioRxiv.

Salay, L.D., Ishiko, N., and Huberman, A.D. (2018). A midline thalamic circuit determines reactions to visual threat. Nature 557, 183–189.

Samsonovich, A., and McNaughton, B.L. (1997). Path integration and cognitive mapping in a continuous attractor neural network model. J. Neurosci. 17, 5900–5920.

Sanders, H., Wilson, M.A., and Gershman, S.J. (2020). Hippocampal remapping as hidden state inference. Elife 9.

Sargolini, F., Fyhn, M., Hafting, T., McNaughton, B.L., Witter, M.P., Moser, M.-B., and Moser, E.I. (2006). Conjunctive representation of position, direction, and velocity in entorhinal cortex. Science 312, 758–762.

Schönemann, P.H. (1966). A generalized solution of the orthogonal procrustes problem. Psychometrika 31, 1–10.

Seely, J.S., Kaufman, M.T., Ryu, S.I., Shenoy, K.V., Cunningham, J.P., and Churchland, M.M. (2016). Tensor analysis reveals distinct population structure that parallels the different computational roles of areas M1 and V1. PLoS Comput. Biol. 12, e1005164.

Seung, H.S. (1996). How the brain keeps the eyes still. Proc. Natl. Acad. Sci. U. S. A. 93, 13339–13344.

Sheintuch, L., Geva, N., Baumer, H., Rechavi, Y., Rubin, A., and Ziv, Y. (2020). Multiple Maps of the Same Spatial Context Can Stably Coexist in the Mouse Hippocampus. Curr. Biol. 30, 1467–1476.e6.

Shlens, J. (2005). A tutorial on Principal Components Analysis. April 7, 2014.

Sikaroudi, A.E., and Park, C. (2019). A mixture of linear-linear regression models for a linear-circular regression. Stat. Modelling 1471082X19881840.

Singh, A.P., and Gordon, G.J. (2008). A Unified View of Matrix Factorization Models. In Machine Learning and Knowledge Discovery in Databases, (Springer Berlin Heidelberg), pp. 358–373.

Skaggs, W.E., Knierim, J.J., Kudrimoti, H.S., and McNaughton, B.L. (1995). A model of the neural basis of the rat’s sense of direction. Adv. Neural Inf. Process. Syst. 7, 173–180.

Skaggs, W.E., McNaughton, B.L., Wilson, M.A., and Barnes, C.A. (1996). Theta phase precession in hippocampal neuronal populations and the compression of temporal sequences. Hippocampus 6, 149–172.

Solstad, T., Boccara, C.N., Kropff, E., Moser, M.-B., and Moser, E.I. (2008). Representation of geometric borders in the entorhinal cortex. Science 322, 1865–1868.

Stensola, H., Stensola, T., Solstad, T., Frøland, K., Moser, M.-B., and Moser, E.I. (2012). The entorhinal grid map is discretized. Nature 492, 72–78.

Stringer, C., Pachitariu, M., Steinmetz, N., Reddy, C.B., Carandini, M., and Harris, K.D. (2019). Spontaneous behaviors drive multidimensional, brainwide activity. Science 364, 255.

Stringer, S.M., Rolls, E.T., and Trappenberg, T.P. (2004). Self-organising continuous attractor networks with multiple activity packets, and the representation of space. Neural Networks 17, 5–27.

Udell, M., Horn, C., Zadeh, R., and Boyd, S. (2016). Generalized Low Rank Models.

Vinck, M., Batista-Brito, R., Knoblich, U., and Cardin, J.A. (2015). Arousal and locomotion make distinct contributions to cortical activity patterns and visual encoding. Neuron 86, 740–754.

Wang, F., and Gelfand, A.E. (2013). Directional data analysis under the general projected normal distribution. Stat. Methodol. 10, 113–127.

Widloski, J., and Fiete, I.R. (2014). A model of grid cell development through spatial exploration and spike time-dependent plasticity. Neuron 83, 481–495.

Williams, A.H., Kim, T.H., Wang, F., Vyas, S., Ryu, S.I., Shenoy, K.V., Schnitzer, M., Kolda, T.G., and Ganguli, S. (2018). Unsupervised Discovery of Demixed, Low-Dimensional Neural Dynamics across Multiple Timescales through Tensor Component Analysis. Neuron 98, 1099–1115.e8.

Wold, S. (1978). Cross-Validatory Estimation of the Number of Components in Factor and Principal Components Models. Technometrics 20, 397–405.

Yoon, K., Buice, M.A., Barry, C., Hayman, R., Burgess, N., and Fiete, I.R. (2013). Specific evidence of low-dimensional continuous attractor dynamics in grid cells. Nat. Neurosci. 16, 1077–1084.

Yu, B.M., Cunningham, J.P., Santhanam, G., Ryu, S.I., Shenoy, K.V., and Sahani, M. (2009). Gaussian-Process Factor Analysis for Low-Dimensional Single-Trial Analysis of Neural Population Activity. Journal of Neurophysiology 102, 614–635.

Zhang, K. (1996). Representation of spatial orientation by the intrinsic dynamics of the head-direction cell ensemble: a theory. J. Neurosci. 16, 2112–2126.

